# Induced ubiquitination bypasses canonical ERAD to drive ER protein degradation

**DOI:** 10.1101/2025.11.28.691080

**Authors:** Sydney J. Tomlinson, Sean L. Johnson, Anna H. Kroskrity, Yun Hu, Kirandeep K. Deol, Cynthia Y. Zhang, Cynthia A. Harris, Daniel K. Nomura, James A. Olzmann

## Abstract

Heterobifunctional proteolysis-targeting chimeras (PROTACs) have emerged as a powerful strategy to degrade disease-relevant proteins, enabling targeting of previously “undruggable” proteins. Current degrader molecules primarily target cytosolic substrates, yet nearly one-third of the proteome resides in or transits the endoplasmic reticulum (ER), including receptors, secreted factors, and biosynthetic enzymes with high therapeutic relevance. Whether ER-localized proteins can be broadly targeted for induced degradation remains an open question. To address this gap, we employed a panel of fluorescent reporter cell lines and used the dTAG chemical-genetic system to recruit cytosolic E3 ligases. While lumenal substrates segregated from the cytosol were resistant to degradation, recruitment of cytosolic ligases effectively degraded ER membrane proteins across multiple topologies and with post-translational modifications. CRISPR genetic screens revealed that the induced degradation required the expected cullin RING ligase complexes but surprisingly bypassed ER-associated degradation (ERAD) machinery, with the exception of the AAA ATPase VCP. Mechanistic studies demonstrated that substrate ubiquitination was essential for VCP binding, and cleavage of ubiquitin chains released VCP, suggesting a model in which VCP directly extracts substrates independent of a dislocation apparatus. Extending this strategy to an endogenous substrate, we synthesized an HMGCR ERAD-TAC by linking atorvastatin to a cereblon E3 ligase recruiter and found that HMGCR degradation was likewise VCP-dependent. Together, these findings demonstrate that ER membrane proteins are generally susceptible to induced degradation via cytosolic ligase recruitment, uncovering a VCP-centered mechanism that operates independently of membrane-embedded ERAD machinery. This work establishes foundational principles for extending targeted protein degradation to the early secretory pathway.

**SIGNIFICANCE STATEMENT:** Targeted protein degradation has transformed drug discovery. Nearly one-third of the proteome reside in or transit the endoplasmic reticulum (ER), a compartment rich in therapeutically relevant but structurally complex targets. Whether these ER proteins can be broadly degraded using PROTACs has remained unknown. Here, we define the minimal requirements for degrading ER membrane proteins by recruiting cytosolic E3 ligases. Using chemical-genetic tools, genetic screens, and a statin-based degrader, we show that ubiquitination engages the VCP extraction machinery, enabling degradation of diverse ER membrane proteins independent of canonical ER-associated degradation components. These findings reveal a ubiquitin-driven route for membrane protein turnover, expand the landscape of druggable ER proteins, and establish principles for designing degraders operating in the early secretory pathway.

## INTRODUCTION

Heterobifunctional small molecule proteolysis-targeting chimeras (PROTACs) have emerged as a powerful therapeutic modality to selectively eliminate disease-driving proteins^1–3^. By simultaneously binding a target protein and an E3 ubiquitin ligase, PROTACs induce their proximity and promote target ubiquitination and subsequent proteasomal degradation^1–3^. Because they act catalytically and do not depend on inhibiting enzymatic activity, PROTACs enable targeting of non-enzymatic or otherwise difficult-to-drug proteins and can retain activity in the face of resistance mutations that diminish inhibitor binding^1–3^. The development of PROTACs has progressed rapidly, with many molecules advancing into clinical trials^1–3^.

Despite this momentum, most degrader development has focused on cytosolic proteins. However, a substantial fraction of the proteome resides in the endoplasmic reticulum (ER), including proteins that regulate lipid metabolism, proteostasis, redox homeostasis, and stress signaling, as well as secreted factors and receptors that transit the ER en route to other cellular destinations^4,5^. Many of these ER-localized or ER-trafficked proteins play key roles in human disease and represent compelling therapeutic targets^5,6^, yet their structural diversity and membrane-embedded environments may render them refractory to conventional small molecule inhibition. Moreover, ER proteins possess a broad range of topologies, including soluble lumenal proteins and integral membrane proteins with single or multiple transmembrane domains, as well as extensive post-translational modifications such as glycosylation and disulfide bonds^5,7,8^. These features raise fundamental questions about the accessibility of ER proteins to PROTAC-mediated ubiquitination and subsequent targeting for proteasomal degradation.

The turnover of ER proteins is mediated by ER-associated degradation (ERAD), a coordinated pathway that includes substrate recognition, ubiquitination, dislocation, and extraction from the membrane by the AAA ATPase VCP (also known as p97), followed by proteasomal degradation^8,9^. Recent reports demonstrate that select multi-pass ER transmembrane proteins, including solute carrier family members^10^ and HMG-CoA reductase (HMGCR)^11,12^, can be degraded following chemical recruitment of cytosolic E3 ligases. These observations suggest that cytosolic ligase recruitment may, under some circumstances, be sufficient to initiate degradation of ER proteins. However, it remains unknown whether such mechanisms are broadly generalizable, what classes of ER proteins are amenable to induced degradation, and what cellular machinery is required to support this process. The extent to which PROTAC-mediated ubiquitination interfaces with ERAD components, and whether distinct extraction or quality-control pathways are engaged, is similarly unresolved.

Here, we systematically evaluate the ability of PROTACs that recruit cytosolic E3 ligases to induce degradation of ER proteins with diverse topologies and post-translational modifications. We find that cytosolic ligase recruitment drives broad clearance of ER membrane proteins through a mechanism that proceeds independently of canonical ERAD components, except for a requirement for the AAA ATPase VCP to extract the ubiquitinated substrates from the membrane. By establishing the mechanistic principles governing cytosolic E3 ligase-driven degradation of ER-localized proteins, this work provides a foundation for expanding targeted protein degradation to a major class of proteins that exhibit unique challenges for existing drug modalities.

## RESULTS

### Fluorescent reporter cell lines to monitor chemically induced degradation of ER proteins

To systematically assess whether ER proteins can be degraded upon recruitment of cytosolic E3 ligases, we generated a panel of GFP-based fluorescent reporter cell lines in U-2 OS and HEK293T backgrounds using the dTAG system^13,14^. In this system, the protein of interest (POI) is fused to FKBP^F36V^, a small 12-kDa protein tag, that can selectively bind a heterobifunctional degrader^13,14^. The small molecule dTAGv-1 and dTAG-13 bind FKBP^F36V^ and recruit either VHL or CRBN-containing cytosolic Cullin E3 ligases, respectively, promoting ubiquitination of the tagged substrate^13,14^ (**Fig 1A**).

**Fig. 1.**
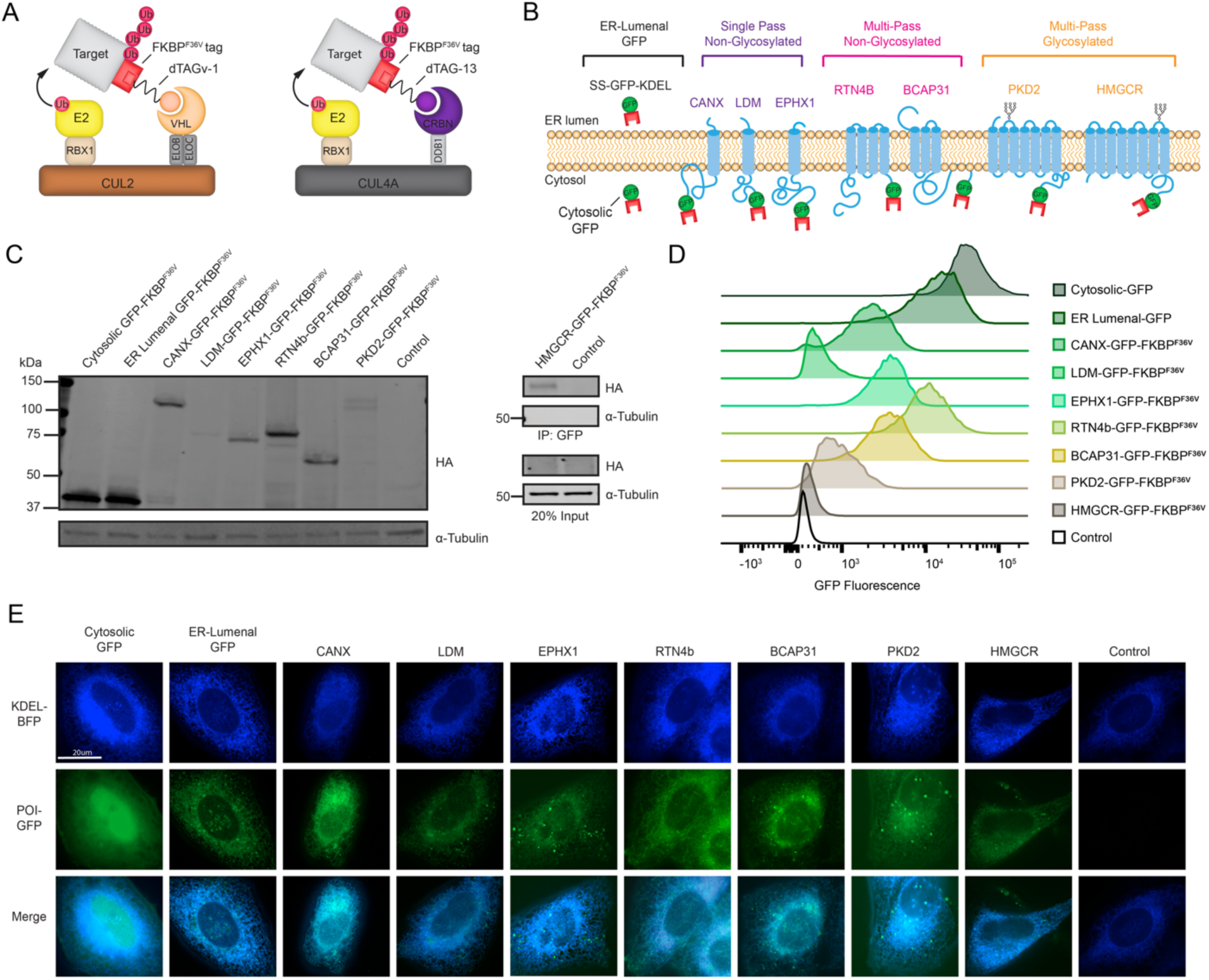
A panel of fluorescent ER protein reporter cell lines for analysis of induced degradation. **(A)** Schematic representation of the dTAG system. dTAGv-1 (orange) and dTAG-13 (purple) bind FKBP^F36V^-tagged proteins and recruit VHL- and CRBN-containing cullin E3 ligase complexes, respectively. **(B)** Diagram of fluorescent protein reporter panel, including ER-lumenal GFP (SS-GFP-KDEL), single-pass non-glycosylated (CANX, LDM, EPHX1), multi-pass non-glycosylated (BCAP31 and RTN4b) and multi-pass glycosylated (PKD2 and HMGCR) reporters. **(C)** Left: Immunoblot of lysates from the indicated U-2 OS reporter cell lines probed for the HA tag and α-tubulin as a loading control. Right: GFP immunoprecipitation (GFP-trap) from wild-type (WT) and HMGCR-GFP-FKBP^F36V^ U-2 OS cells followed by HA immunoblotting; a 20% input lysate is also analyzed. **(D)** Flow cytometry histograms of GFP fluorescence in wild-type U-2 OS and fluorescent reporter cell lines. **(E)** Fluorescence microscopy of GFP-tagged proteins (green) with the ER marker BFP-KDEL (blue). Scale bar, 20 μm.

We selected a panel of ER-resident substrates that share a cytosolic-facing C-terminus to ensure FKBP^F36V^ accessibility to the recruited ubiquitination machinery, and that together reflect the topological and post-translational diversity of ER proteins. Substrates were C-terminally tagged with GFP-FKBP^F36V^-3×HA and stably expressed in both U-2 OS and HEK293T cells. The panel included single-pass, non-glycosylated proteins (calnexin [CANX], lanosterol 14α-demethylase [LDM], and epoxide hydrolase 1 [EPHX1]); multi-pass, non-glycosylated proteins (Reticulon-4b [RTN4b] and BCAP31); and multi-pass, glycosylated substrates (polycystin-2 [PKD2] and the polytopic transmembrane region of HMG-CoA reductase [HMGCR]) (**Fig. 1B**). In addition, we generated an ER lumenal GFP-FKBP^F36V^-3×HA reporter and a cytosolic GFP-FKBP^F36V^-3×HA reporter

Immunoblot analysis confirmed that the reporter substrates were expressed at their expected molecular weight (**Fig. 1C**). Because HMGCR-GFP-FKBP^F36V^ is present at low basal abundance, we performed a GFP immunoprecipitation to enrich the protein and verify its size (**Fig. 1C**). Flow cytometry demonstrated GFP expression as a single peak in each reporter line (**Fig. 1D**), and fluorescence imaging showed ER-localized signal for the ER lumenal and transmembrane substrates, whereas the cytosolic GFP-FKBP^F36V^ protein was diffusely distributed throughout the cytosol and nucleoplasm (**Fig. 1E, Supplementary Fig. S1**).

Together, these data establish a panel of fluorescence-based reporter cell lines for monitoring ER protein abundance and evaluating the effects of chemically induced recruitment of cytosolic E3 ligases.

### E3 Ligase Recruitment Enables Clearance of Diverse ER Transmembrane Proteins

Immunoblotting using anti-HA and anti-GFP antibodies revealed that increasing doses of dTAG-v1 induced the degradation of representative ER transmembrane (ER-TM) proteins across topological and glycosylation classes, including single-pass non-glycosylated (CANX-GFP-FKBP^F36V^), multi-pass non-glycosylated (BCAP31-GFP-FKBP^F36V^), and multi-pass glycosylated reporters (PKD2-GFP-FKBP^F36V^) (**Fig. 2A**). As expected, the cytosolic GFP-FKBP^F36V^ control was efficiently degraded (**Fig. 2A**). In contrast, the ER-lumenal GFP-FKBP^F36V^ reporter remained stable (**Fig. 2A**), consistent with its inaccessibility to cytosolic E3 ligase recruitment due to spatial segregation in distinct subcellular compartments. Fluorescence microscopy confirmed the loss of ER-localized GFP signal upon degrader treatment (**Fig. 2B**), and additional substrates showing comparable behavior (**Supplementary Fig. S2A,B**). Flow cytometry analyses of the complete panel similarly demonstrated a concentration-dependent loss of GFP signal upon dTAG-v1 or dTAG-13 treatment for cytosolic GFP and ER-TM substrates, but not the ER lumenal substrate (**Fig. 2C**). To determine whether degradation of ER-TM proteins by cytosolic E3 ligase recruitment depends on the presence of the GFP tag, we generated U-2 OS reporter lines expressing CANX–FKBP^F36V^ lacking GFP. Dose response analysis with dTAG-v1 revealed degradation kinetics and efficiency comparable to the GFP-tagged CANX-GFP-FKBP^F36V^ construct, indicating that the GFP tag is not required for substrate recognition or degradation (**Supplementary Fig. S2C**). In addition, parallel analyses in HEK293T cells yielded similar degradation profiles (**Supplementary Fig. S3A,B**), indicating that cytosolic E3 ligand recruitment drives ER-TM protein degradation in a cell line-independent manner.

**Fig. 2.**
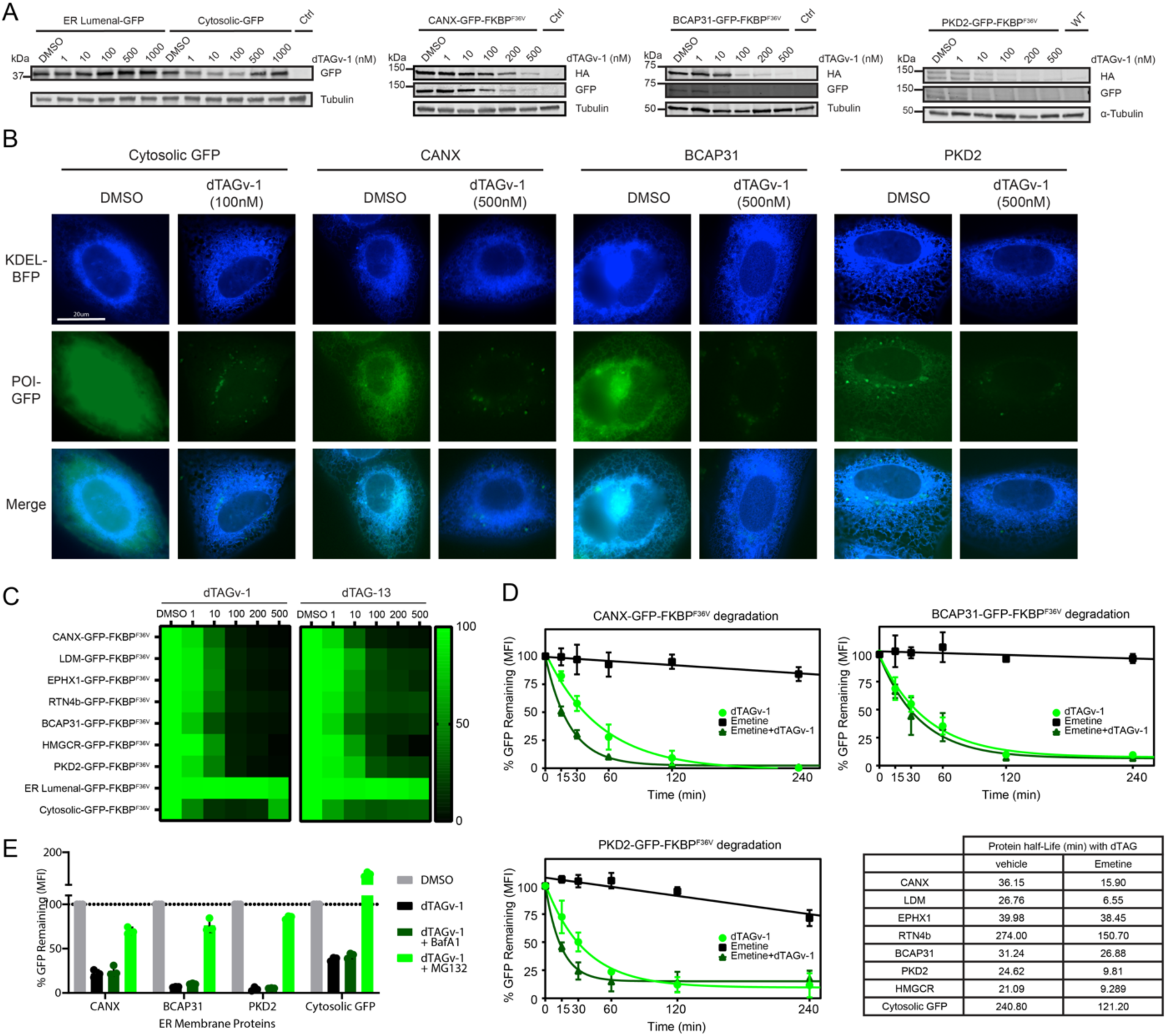
Recruitment of cytosolic E3 ligases induces rapid degradation of diverse ER-TM proteins. ER-TM proteins from each topological and glycosylation classes were analyzed, including single-pass non-glycosylated (CANX-GFP-FKBP^F36V^), multi-pass non-glycosylated (BCAP31-GFP-FKBP^F36V^), and multi-pass glycosylated reporters (PKD2-GFP-FKBP^F36V^) along with positive (cytosolic GFP-FKBP^F36V^) and negative (ER-lumenal GFP-FKBP^F36V^) controls. **(A)** Cells expressing the indicated proteins were treated with the increases concentrations of dTAGv-1 for 24 hr, and the lysates immunoblotted for HA, GFP and α-tubulin. **(B)** Reporter cell lines transiently expressing the ER marker BFP-KDEL (blue) were treated with the indicated concentrations of dTAGv-1 for 24 hr and imaged by fluorescence microscopy. GFP fluorescence (green) indicates the cytosolic and ER-TM reporters. Scale bar, 20 µm. **(C)** Cells were treated with the indicated concentrations of dTAGv-1 or dTAG-13 for 24 hr. GFP fluorescence, reflecting reporter levels, was measured by flow cytometry, and the percentage of GFP remaining visualized as a heatmap. **(D)** Cells were treated with 75 µM emetine, 100 nM dTAGv-1 (for cytosolic GFP-FKBPF36V), 500 nM dTAGv-1 (for all other ER-TM proteins), or combinations thereof for the indicated times. GFP fluorescence was quantified by flow cytometry. The inset table shows reporter protein half-lives under vehicle or emetine-treated conditions. **(E)** Cells were pre-treated for 3 hr with either 10 µM MG132 or 250 nM Bafilomycin A1 followed by co-treatment with the same inhibitor and dTAGv-1, or dTAGv-1 alone, at indicated times and doses. GFP fluorescence was measure with flow cytometry.

To measure ER-TM protein degradation kinetics, we performed chase assays in the presence of the translation inhibitor emetine. The addition of dTAG-v1 and emetine resulted in rapid degradation of ER-TM substrates, with half-lives of <40 minutes for most substrates and none exceeding ∼2.5 hours (**Fig. 2C, Supplementary Fig. S4**). Under certain conditions ER proteins are targeted for lysosomal degradation^15^ and PROTAC-mediated degradation of plasma membrane proteins induces endolysosomal trafficking^16^. To determine whether dTAG-induced degradation proceeds through the proteasome or lysosome targeting, cells were treated with the proteasome inhibitor MG132 or the lysosomal inhibitor BafA1. MG132 markedly attenuated dTAG-v1-induced degradation, whereas BafA1 had no effect, indicating that cytosolic E3 ligase recruitment drives proteasome-dependent degradation of ER-TM proteins (**Fig. 2E**).

Collectively, these results demonstrate that recruitment of either VHL- or CRBN-containing cytosolic E3 ligase complexes is sufficient to trigger rapid proteasomal degradation of topologically diverse and glycosylated ER transmembrane proteins.

### Genetic Screens Reveal Factors that Mediate the Induced Degradation of ER-TM Proteins

ER-TM proteins are typically degraded through ERAD, a process in which membrane-embedded complexes recognize misfolded or regulated substrates, ubiquitinate them, and recruit the extraction machinery required for dislocation into the cytosol for proteasomal degradation^5,8,9^. However, how ER-TM proteins that are ubiquitinated directly by cytosolic E3 ligases are dislocated and extracted from the membrane remains unknown. For example, one possibility is that ubiquitination may be sufficient to trigger engagement with ERAD factors that containing ubiquitin binding domains (e.g., UBXD8, UBXD2, or AUP1), which are part of larger membrane-embedded dislocation complexes^5,8,9,17^, to promote extraction.

To identify factors required for the degradation of ER-TM proteins by recruited cytosolic E3 ligases, we performed FACS-based CRISPR-Cas9 loss-of-function screens in three representative reporter cell lines expressing either the cytosolic GFP-FKBP^F36V^, single-pass ER-TM protein CANX-GFP-FKBP^F36V^, and multi-pass glycosylated ER-TM protein PKD2-GFP-FKBP^F36V^. We used a degradation-focused sgRNA library, which targets ∼2,150 genes across the <u>Ub</u>iquitin-proteasome, <u>a</u>utophagy–lysosome, and <u>ER</u>AD (UBALER) pathways. This library was generated by adding sgRNAs targeting ∼150 additional genes encoding glycosylation enzymes, ERAD complexes, and dislocation/retrotranslocation factors to our previously generated UBAL sgRNA library^18^. Cas9-expressing reporter cell lines were transduced with the UBALER library and treated with dTAG-v1 until approximately 20% of cells retained GFP signal, after which the brightest 5% (GFP^high^) and dimmest 75% (GFP^low^) cells were isolated by FACS for sequencing. Gene enrichment scores and confidence values were calculated using the Cas9 High-Throughput Maximum Likelihood Estimator (casTLE) framework^19–21^ (**Fig. 3A,B and Supplementary Table S1**). Genes enriched in the GFP^high^ population relative to the GFP^low^ population represent loss-of-function events that stabilize the reporter and impair induced degradation. The screens maintained excellent library representation (**Supplementary Fig. S5A**) and together enabled systematic comparison of pathways that mediate cytosolic E3 ligase-induced degradation across ER-TM proteins of differing topologies and complexity (**Fig. 3A-D**).

**Fig. 3.**
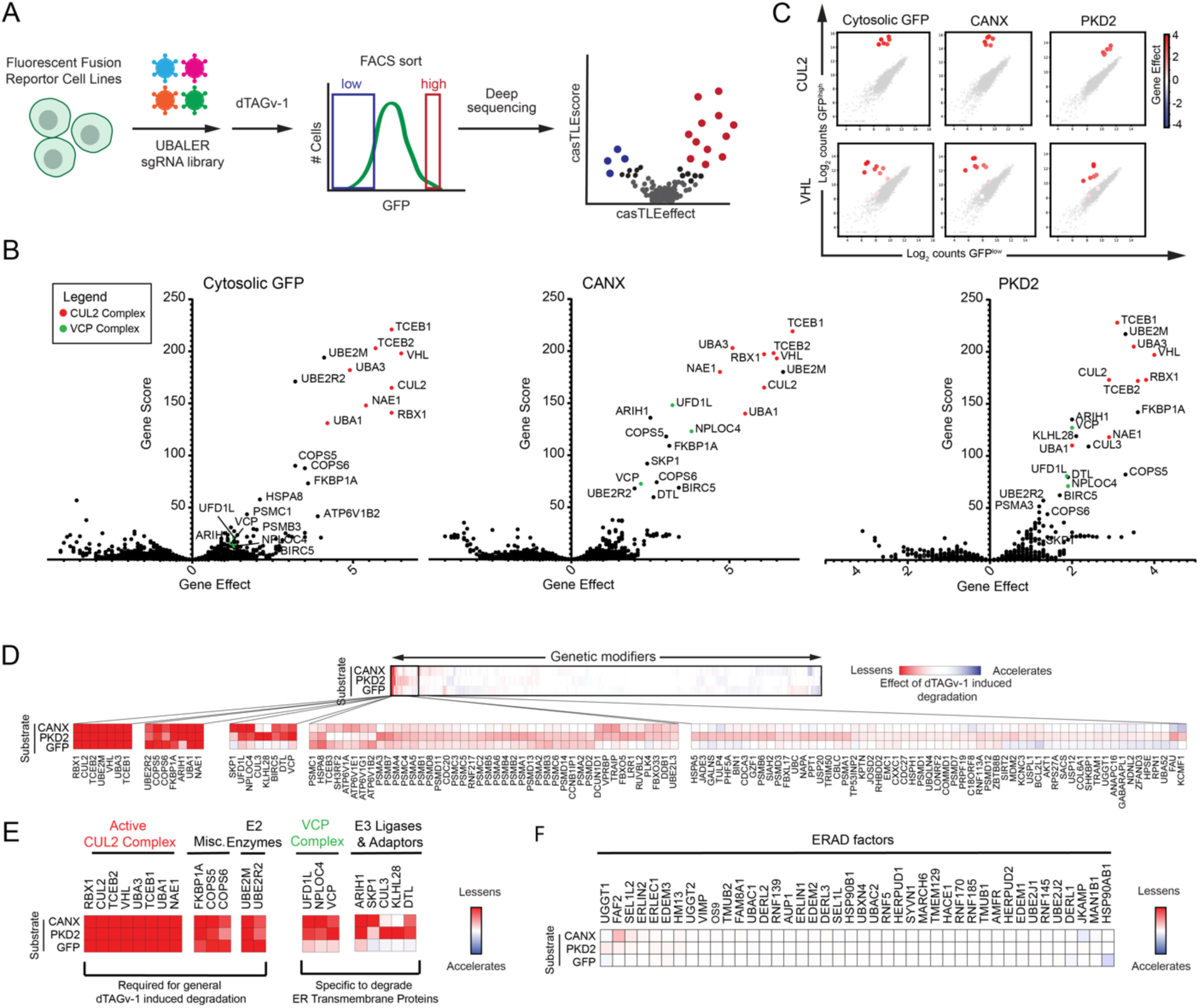
CRISPR-Cas9 screens reveal the mechanism of ER-TM protein induced degradation. **(A)** Schematic of the UBALER CRISPR–Cas9 screening strategy. Fluorescent ER–TM reporter cell lines were treated with dTAGv-1 and sorted by FACS into high- and low-GFP populations. sgRNA representation in each population was quantified by deep sequencing to calculate gene effects and confidence scores (gene scores). **(B)** Scatter plots of gene effect versus gene score for cytosolic GFP, CANX, and PKD2 reporter screens. Genes encoding components of the CUL2 complex (red) and the VCP complex (green) are highlighted. **(C)** Cloud plots indicating count numbers corresponding to *CUL2* and *VHL* (color scale) and control (gray scale) sgRNAs for each screen. **(D)** Heatmap of gene scores across all reporter lines with unbiased hierarchical clustering. Genes whose knockout lessens dTAGv-1-induced degradation are shown in red; those that accelerate degradation are shown in blue. **(E)** Functional clustering of top hits. **(F)** Heatmap of ERAD-related genes.

Across all three reporter backgrounds, the CUL2-VHL E3 ligase complex (*CUL2, VHL, TCEB1, TCEB2, RBX1*) and the neddylation machinery (*UBA3, NAE1*) emerged as the strongest hits, consistent with their essential role in dTAG-v1-induced degradation. For both VHL and CUL2, at least 8 of 10 sgRNAs were consistently enriched in GFP^high^ cells and depleted in GFP^low^ cells across all screens, driving the high confidence for these factors (**Fig. 3C, Supplementary Fig. S5B-D**). As dTAG-v1 functions by recruiting the VHL-containing CUL2 complex, these results were expected and confirm the fidelity of the induced degradation system and our genetic screening approach.

Among the next highest-scoring genes were *VCP* and its cofactors *UFD1L* and *NPLOC4*, which together form the VCP-UFD1-NPL4 extraction complex (**Fig. 3D,E, Supplementary Fig. S5B-D**). This is notable because VCP is the AAA ATPase responsible for substrate extraction from the ER membrane. Strikingly, no other canonical ERAD components, including substrate recognition factors, glycan-dependent lectins, or the membrane-embedded dislocation complexes (e.g., HRD1/SEL1L or derlins), were significantly enriched in any screen (**Fig. 3F**). This suggests that cytosolic E3 ligase-induced degradation of ER-TM proteins proceeds independently of the canonical ERAD recognition machinery and relies primarily on VCP for membrane extraction without the need for a membrane-embedded dislocation channel, even for the glycosylated polytopic membrane protein PKD2. Together, these genetic screens identify a minimal set of factors that mediate induced degradation of ER-TM proteins, engagement of a cytosolic E3 ligase and extraction by VCP, without a requirement for upstream ERAD substrate processing.

### VCP Is Essential for Cytosolic E3 Ligase-Induced ER Protein Extraction and Degradation

The identification of VCP and its cofactors in our genetic screens suggests that VCP is required to extract ER-TM proteins during cytosolic E3 ligase-mediated degradation. To directly test this, we performed degrader time-course assays in the presence of two mechanistically distinct VCP inhibitors, CB-5083^22,23^ and NMS-873^24^. Inhibition of VCP markedly attenuated dTAG-v1-induced degradation across all ER-TM substrates (**Fig. 4A, Supplementary Fig. S6A**), similar to the effect of the neddylation inhibitor MLN4924, which blocks activation of the CUL2 ligase complex. This requirement for VCP was not specific to CUL2 recruitment, as VCP inhibition also attenuated dTAG-13-induced degradation, which recruits a distinct E3 ligase complex (i.e., CUL4A-CRBN) (**Fig. 4B**). Immunoblot analysis also confirmed the attenuation of ER-TM protein degradation in response to VCP inhibition (**Fig. 4C**).

**Fig. 4.**
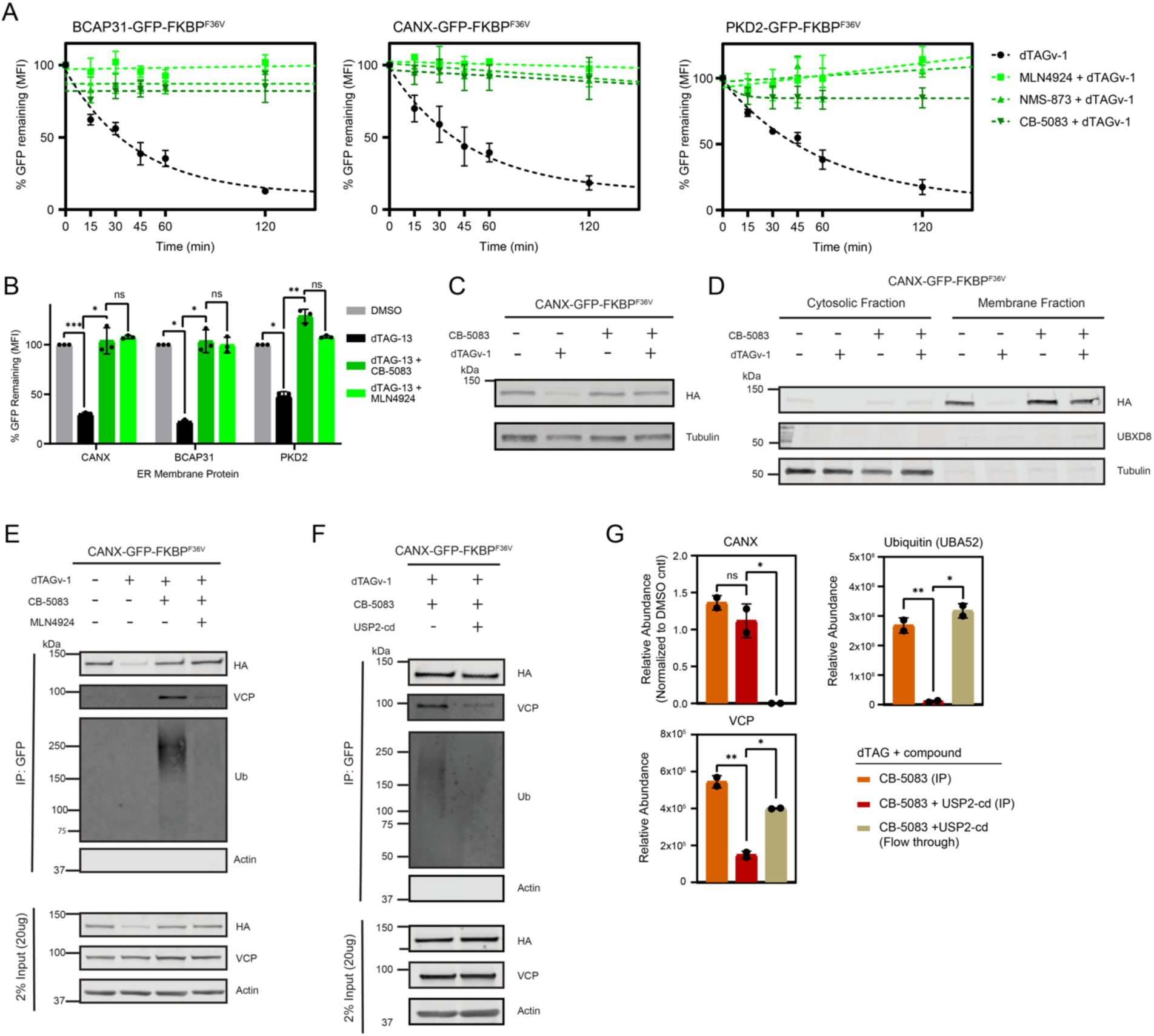
Induced ER-TM protein degradation requires ubiquitin-dependent VCP-mediated extraction. Specified U-2 OS fluorescent ER-TM protein reporter cell lines were pre-treated for 30 min with 5 µM CB-5083, 25 µM NMS-873, or 500 nM MLN4924, followed by co-treatment with 500 nM dTAGv-1 or dTAG-13 for the indicated times unless otherwise noted; DMSO served as the vehicle control. **(A-B)** Kinetics and quantification of GFP fluorescence decay in the indicated reporter cell lines. **(C)** Immunoblot of CANX-GFP-FKBP^F36V^ cell lysates treated with as indicated and probed for HA-tagged protein and α-tubulin. **(D)** Cytosolic and membrane fractions collected from cells treated as indicated and immunoblotted for HA, UBXD8 (membrane marker), and α-tubulin (cytosolic marker). **(E)** GFP immunoprecipitation (IP) from cells treated as indicated followed by immunoblotting for HA, VCP, ubiquitin, and β-actin (n=1). **(F)** GFP-IP samples from cells treated with CB-5083 and dTAGv-1, incubated with the USP2 catalytic domain (USP2-cd) where indicated to remove ubiquitin chains, and immunoblotted for HA, VCP, ubiquitin, and β-actin. **(G)** Quantitative proteomic analysis of GFP-IP samples from **(F)** showing relative abundances.

Consistent with VCP’s established role in ER membrane protein extraction, inhibition of VCP resulted in the accumulation of CANX-GFP-FKBP^F36V^ in the ER-enriched membrane fraction rather than in the cytosol (**Fig. 4D**). This dependence on VCP was also observed in the cell line expressing CANX-FKBP^F36V^ lacking the GFP tag, confirming that VCP function is required independent of the GFP tag (**Supplementary Fig. S6B**).

Together, these results demonstrate that VCP activity is required to extract ER-TM proteins from the membrane during degradation induced by cytosolic E3 ligase recruitment.

### Ubiquitination Initiates and Sustains VCP Binding During Induced ER Membrane Protein Extraction

To assess the role of ubiquitination in VCP engagement during cytosolic E3 ligase-mediated degradation, we first examined VCP association with ER-TM substrates by immunoprecipitation. Inhibition of VCP with CB-5083 trapped VCP in complex with ubiquitinated ER-TM proteins during degrader treatment (**Fig. 4E, Supplementary Fig. S6C**). However, when cullin-mediated ubiquitination was blocked using the neddylation inhibitor MLN4924, both ubiquitination and VCP interaction were lost (**Fig. 4E, Supplementary Fig. S6C**). To test whether ubiquitination is required to maintain VCP association, ER-TM proteins were immunoprecipitated and treated on-bead with the deubiquitinase USP2. Removal of ubiquitin abolished VCP binding (**Fig. 4F**). Proteomic analysis of these samples confirmed the loss of both ubiquitin and VCP from the immunoprecipitated fraction and their corresponding presence in the flow-through (**Fig. 4G**). These data indicate that ubiquitination is required to initiate and maintain VCP association with ER-TM substrates during induced degradation.

We next asked whether VCP recruitment to ER-TM proteins is sufficient to trigger degradation. To test this possibility, we stably expressed CANX-GFP-ABI1 with effectors fused to a GFP nanobody (vhhGFP), promoting constitutive substrate-effector binding. While CRBN-vhhGFP drove robust degradation of CANX-GFP-ABI1, VCP-, UFD1, and NPLOC4–vhhGFP fusions induced only minimal loss of the reporter (**Supplementary Fig. S7A-D**). Thus, recruitment of VCP or its cofactors alone is insufficient to promote ER-TM degradation.

Together, these results demonstrate that ubiquitination is required to initiate and maintain VCP engagement, enabling extraction of ER-TM substrates during cytosolic E3 ligase-induced degradation, whereas VCP recruitment without ubiquitination is insufficient to drive turnover.

### A Targeted ERAD-Based Degrader Counteracts Statin-Induced HMGCR Accumulation

Our findings using the chemogenetic dTAG system indicate that ubiquitination drives VCP-dependent extraction and degradation of ER-TM proteins following cytosolic E3 ligase recruitment. To test whether this mechanism extends to an endogenous ER-resident substrate, we synthesized an HMGCR ERAD-TAC (11j/P22A^12^), in which atorvastatin is linked to the CRBN E3 ligase recruiter pomalidomide (**Fig. 5A**). HMGCR provides a relevant biological context because, although statins effectively inhibit its catalytic activity and reduce cholesterol biosynthesis they also diminish sterol-dependent HMGCR turnover and activate SREBP-dependent transcription of *HMGCR*^25–29^. This leads to a compensatory increase in HMGCR protein abundance, a feedback response that contributes to statin resistance^25–29^. Consistent with its feedback regulation, atorvastatin treatment of sterol-depleted Huh7 cells elicited a robust, dose-dependent accumulation of HMGCR protein (**Fig. 5B)**. We therefore asked whether 11j/P22A could prevent this compensatory stabilization during statin treatment. Remarkably, 11j/P22A dose-dependently attenuated this compensatory accumulation (**Fig. 5C**), with a measurable reduction as early as 4 hours of treatment (**Fig. 5D**). Because 11j/P22A contains an atorvastatin moiety that inhibits HMGCR enzymatic activity, we anticipated activation of SREBP-dependent transcription. Indeed, both 11j/P22A and atorvastatin increased mRNA levels of the canonical SREBP2 targets *Hmgcr*, *Ldlr*, and *Fasn*, consistent with 11j/P22A binding and inhibition of HMGCR (**Fig. 5E**). Thus, the statin-based HMGCR degrader maintains enzymatic inhibition while also preventing the compensatory accumulation of HMGCR.

**Fig. 5.**
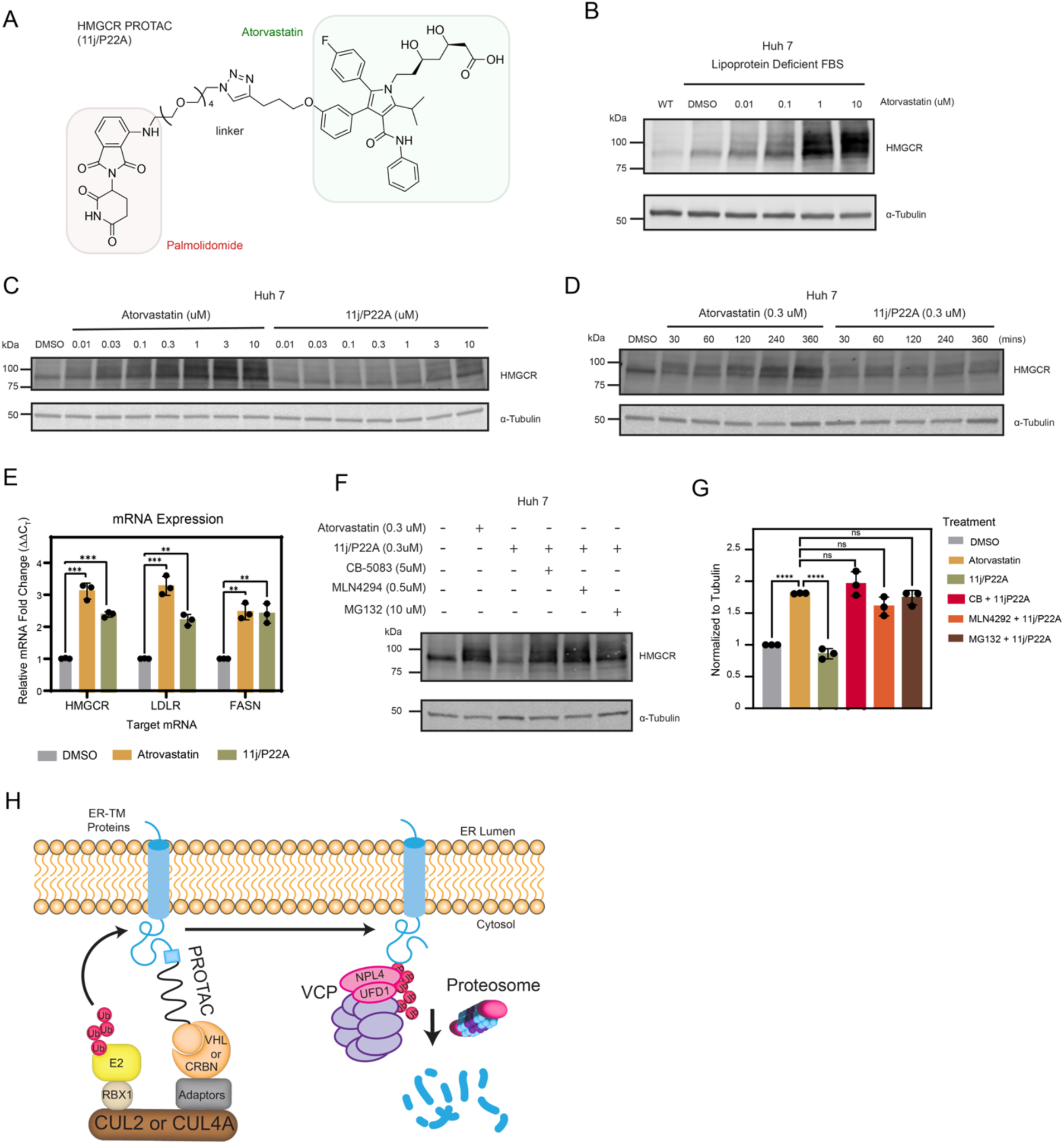
VCP is required for endogenous HMGCR degradation by a CRBN-based PROTAC. **(A)** Chemical structure the endogenous HMGCR ERAD-TAC (11j/P22A^12^). Huh7 cells were cultured for 16 hr in medium containing 10% lipoprotein-deficient FBS prior to treatment. Cells were then treated with atorvastatin or 11j/P22A at the indicated concentrations and times, as specified for each panel. For inhibitor studies, cells were pre-treated for 30 min with 5 µM CB-5083, 0.5 µM MLN4924, or 10 µM MG132 (3hr pre-treatment) followed by co-treatment with the same inhibitor and 0.3 µM 11j/P22A or atorvastatin. **(B-D)** Immunoblots showing HMGCR protein abundance following treatment with atorvastatin or 11j/P22A at the indicated concentrations and times. **(E)** Relative *HMGCR, LDLR, and FASN* mRNA fold change after 24 h treatment with 0.3 µM atorvastatin or 0.3 µM 11j/P22A. **(G)** Quantification of HMGCR protein levels from immunoblots in **(F)**. **(H)** Proposed mechanism for ER-TM protein degradation by recruitment of cytosolic E3 ligase, in which the ubiquitination of the ER protein results in the recruitment of the VCP-UFD1-NPL4 complex. VCP directly binds ubiquitin, mediate initiation of unfolding and substrate processing, and extracts the substrate from the membrane independently of a membrane-embedded dislocation channel.

### VCP Is Required for the Induced Degradation of Endogenous HMGCR

To assess whether the degradation of endogenous HMGCR by the CRBN-based PROTAC requires VCP, sterol-depleted Huh7 cells were treated with atorvastatin alone, 11j/P22A alone, or 11j/P22A in combination with inhibitors of VCP (CB-5083), the proteasome (MG132), or cullin activation (MLN4924) (**Fig. 5F,G**). As expected, atorvastatin alone increased HMGCR abundance, whereas 11j/P22A reduced HMGCR levels. Co-treatment of 11j/P22A with CB-5083 or MG132 led to an increase in HMGCR to levels comparable to atorvastatin alone (**Fig. 5F,G**), demonstrating that 11j/P22A-mediated degradation requires both VCP activity and the proteasome. Likewise, MLN4924 fully blocked 11j/P22A-induced HMGCR turnover (**Fig. 5F,G**), confirming that degradation depends on cullin E3 ligase activation. Notably, this indicates that HMGCR loss is driven by the recruited CRBN cullin E3 ligase rather than endogenous ERAD pathways, as HMGCR is normally ubiquitinated by gp78 and TRC8^27,30^, which do not require neddylation for activation.

## DISCUSSION

Together, our findings support a model (**Fig. 5H**) in which PROTAC-mediated recruitment of a cytosolic cullin-based E3 ligase complex (e.g., VHL or CRBN) triggers ubiquitination of ER-TM substrates. The resulting polyubiquitin chains directly recruit the VCP-UFD1-NPL4 complex to drive VCP-dependent extraction of the substrate from the ER membrane, enabling its subsequent proteasomal degradation. By examining a panel of ER-TM proteins spanning diverse topologies and post-translational modifications, we demonstrate that induced degradation can be broadly achieved as long as the substrate presents a cytosolic surface accessible for degrader binding and ubiquitin conjugation. These results indicate that the topological and biophysical constraints of the ER membrane do not inherently limit degrader activity and establish a generalizable framework for targeted protein degradation at the ER.

Our genetic screens highlight the VCP-UFD1-NPL4 complex as a central mediator of ER-TM protein induced degradation. Surprisingly, canonical ERAD factors that mediate retrotranslocation and membrane remodeling, including Hrd1 / Derlin complexes and VCP-interacting adaptors with UBX or UBA domains, were not required. This suggests that induced degradation employing cytosolic E3 ligases can proceed independently of a canonical ERAD dislocon and instead relies on ubiquitination as the primary determinant for VCP recruitment. Consistent with this, both cullin inhibition and on-bead deubiquitination disrupted VCP-substrate interaction, whereas enforced recruitment of VCP or its cofactors in the absence of ubiquitination was insufficient to promote degradation. Thus, ubiquitination is both the initiating and sustaining signal for extraction. Our findings align with previous studies in which viral proteins induce the degradation of ER-resident substrates by recruiting a cytosolic E3 ligase, such as the viral protein VPU recruitment of SCF^β-TrCP^ E3 ligase for the degradation of CD4^31–33^ and the viral protein UL49.5 recruitment of the CUL2-KLHDC3 E3 ligase for the degradation of Transporter Associated with Antigen Processing (TAP)^34^. As in our system, the degradation of CD4 and TAP required the VCP complex, but do not appear to require other classical ERAD dislocation and ubiquitination factors^34^. These findings parallel our conclusions and reinforce the concept that polyubiquitin alone may be sufficient to initiate and direct the extraction of many ER membrane proteins by VCP together with its cofactors^35^. This model is also consistent with findings that the yeast homolog of VCP (i.e., Cdc48) binds primarily to the polyubiquitin chain, rather than the substrate, and uses ubiquitin unfolding to initiate substrate processing and extraction^36–38^. We recognized the limitations of our genetic screens, which leave open the possibility of compensatory pathways or components. However, the emerging findings lead us to speculate that although a dedicated dislocon (i.e., membrane embedded, transmembrane channel) may be essential primarily for lumenal ERAD substrates, which must be threaded across the bilayer, many cytosol-facing ER-TM proteins may be extracted from the membrane independently of a dislocon once sufficiently ubiquitinated.

Finally, we applied this mechanistic framework to the endogenous ER protein HMGCR, the pharmacological target of statins. A CRBN-based statin ERAD-TAC^12^ prevented the compensatory accumulation of HMGCR that typically accompanies statin treatment, and the induced degradation of HMGCR required both VCP activity and the proteasome. These results support the feasibility of an HMGCR degrader as a therapeutic strategy and demonstrate that recruited cullin E3 ligases can substitute for native ER-resident ligases to drive VCP-dependent clearance of endogenous ER-TM proteins. More broadly, this illustrates the translational potential of exploiting this minimal, ubiquitin- and VCP-dependent degradation pathway to modulate membrane protein homeostasis in disease.

In conclusion, our findings define the minimal requirements for cytosolic E3 ligase-mediated degradation of ER membrane proteins and establish a conceptual framework for designing PROTACs and other induced-proximity degraders that harness VCP-dependent extraction. By bridging principles from protein quality control, membrane biology, and targeted protein degradation, this work expands the accessible landscape of druggable membrane proteins and provides a foundation for therapeutic strategies targeting ER-resident disease drivers.

## AUTHOR INFORMATION

Correspondence and requests for materials should be addressed to J.A.O. and D.K.N.

## ACKNOWLEDGEMENTS

This work was also supported by Novartis Biomedical Research and a grant from the National Institutes of Health to J.A.O. (R01DK128099). S.J.T. was supported in part by a University of California Cancer Research Coordinating Committee (CRCC) predoctoral fellowship award. We thank the team of scientist at Novartis for their valuable suggestions and guidance throughout this project. We also thank Dr. Steve Eyles (UMass Amherst, RRID: SCR_019063) for assistance with high-resolution MS acquired on an Orbitrap Fusion mass spectrometer (NIH grant: 1S10OD010645-01A1).

## AUTHOR CONTRIBUTIONS

S.J.T., J.A.O., and D.K.N. conceived of the project, designed the experiments, and wrote the majority of the manuscript. All authors read, edited, and contributed to the manuscript. S.J.T. performed the majority of the experiments with assistance and discussion from A.K. and C.Y.Z. S.L.J. assisted in the cloning and generation of reporter cell lines. K.K.D assisted with proteomics, and along with C.A.H provided important intellectual contributions. Y.H. synthesized the 11j/P22A.

## METHODS

All reagent catalog numbers can be found in **Supplementary Table S2**.

### Plasmids

The gateway compatible destination plasmid, c-terminal GFP-dTAG, was generated by fusing GFP to the n-terminus of FKBPF36V in the pLEX_305-C-dTAG plasmid (Addgene #91798) using megaprimer. Briefly, to introduce the GFP insert, primers containing 20-bp overlaps with insert site in the pLEX_305-C-dTAG plasmid were used to amplify the GFP from a codon-optimized GFP-P2A-BFP gBlock (Integrated DNA Technologies). This insert was then cloned in frame to the c-terminus of the attR2 site of the pLEX_305-C-dTAG plasmid using restriction enzyme-independent fragment insertion by megaprimer cloning. Transformation of this c-terminal GFP-dTAG plasmid was performed in ccdB resistance cells (Invitrogen #A10460).

Codon-optimized gBlocks encoding ER Lumenal-GFP-FKBP^F36V^ (including an ER signal sequence, GFP, and a KDEL retention sequence), Cytosolic-GFP-FKBP^F36V^, and HMGCR-GFP-FKBP^F36V^ were first cloned into the pDONR221 vector using a BP clonase rxn (Invitrogen #11789020), according to the manufacturer’s instructions. These along with the following entry plasmids CANX, LDM (CYP51A1), EPHX1, RTN4b, PKD2, where then cloned into the GFP-dTAG destination plasmid and CANX was also cloned into the pLEX_305-C-dTAG using a LR clonase rxn (Invitrogen #11791020) according to manufacturer’s instructions, and virally transduced into U 2 O-S and HEK293T WT cells (as described in ‘Generation of Membrane Protein-GFP-FKBP^F36V^).

The original pLX301-GFP-ABI1-IRES-BFP plasmid was gifted from the Taipale lab^39^. Gateway cloning was used to generate a ccDB-selectable pLX301-ccDB-ABI1-IRES-BFP gateway-compatible plasmid followed by an additional gateway reaction with a CANX-GFP gene block for the final pLX301-CANX-GFP-ABI1-BFP plasmid. An additional gateway-compatible plasmid provided by the Taipale lab (pLX301-ccDB-vhhGFP) was used to tag VCP and its cofactors with a GFP-targeting nanobody tag that would induce proximity with the GFP-tagged CANX target. Gateway cloning reactions between effector entry plasmids and the pLX301-ccDB-vhhGFP destination vector produced Effector-vhhGFP-containing lentiviral plasmids. U 2 O-S WT cell were virally transduced first with pLX301-CANX-GFP-ABI1-BFP and followed by each Effector-vhhGFP-containing lentiviral plasmid resulting in 5 unique cell lines expression both CANX-GFP and either VCP-vhhGFP, NPLOC4-vhhGFP, UFD1L-vhhGFP, CRBN-vhhGFP, or VHL-vhhGFP (as described in ‘Generation of Membrane Protein-GFP-FKBP^F36V^’).

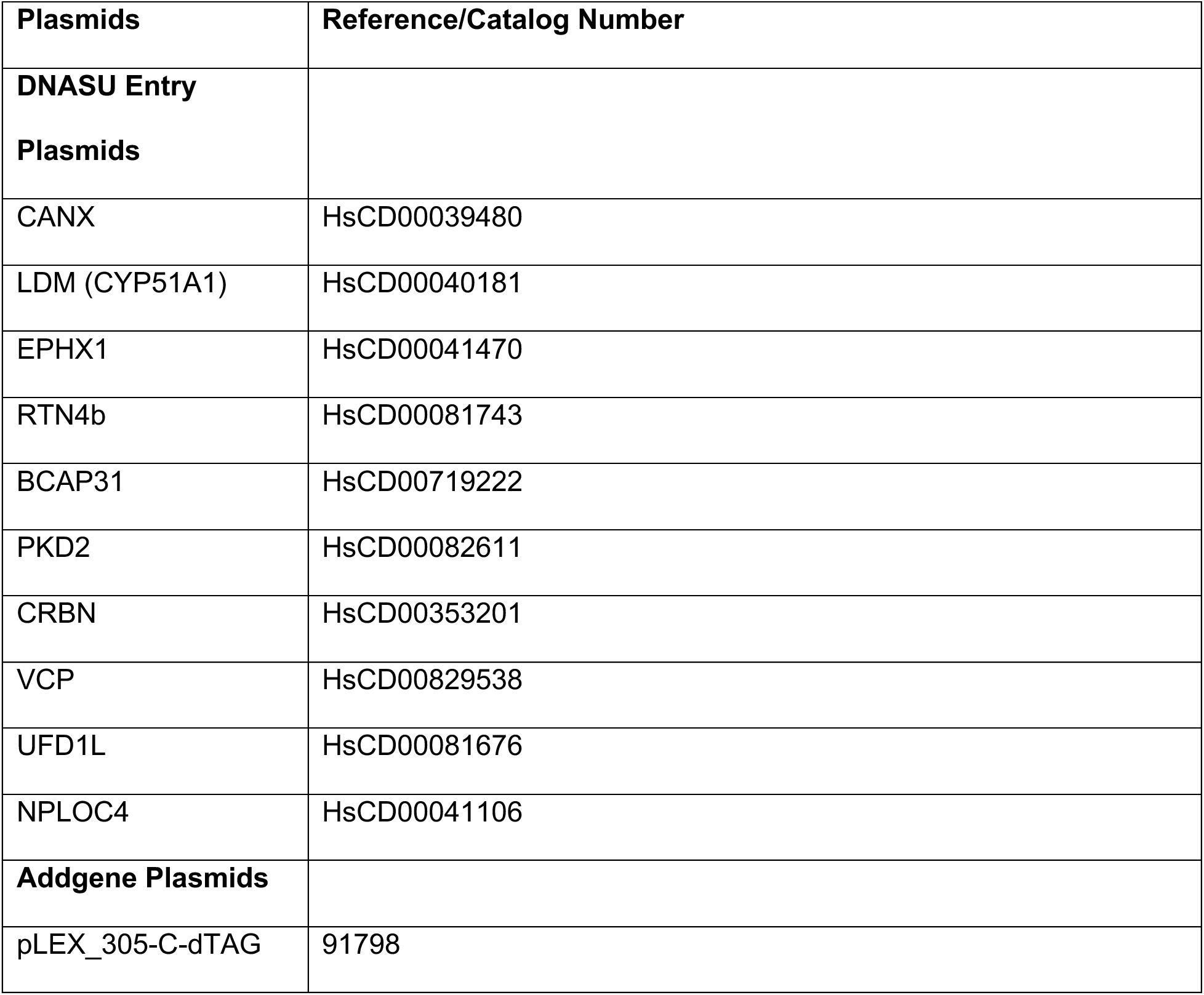
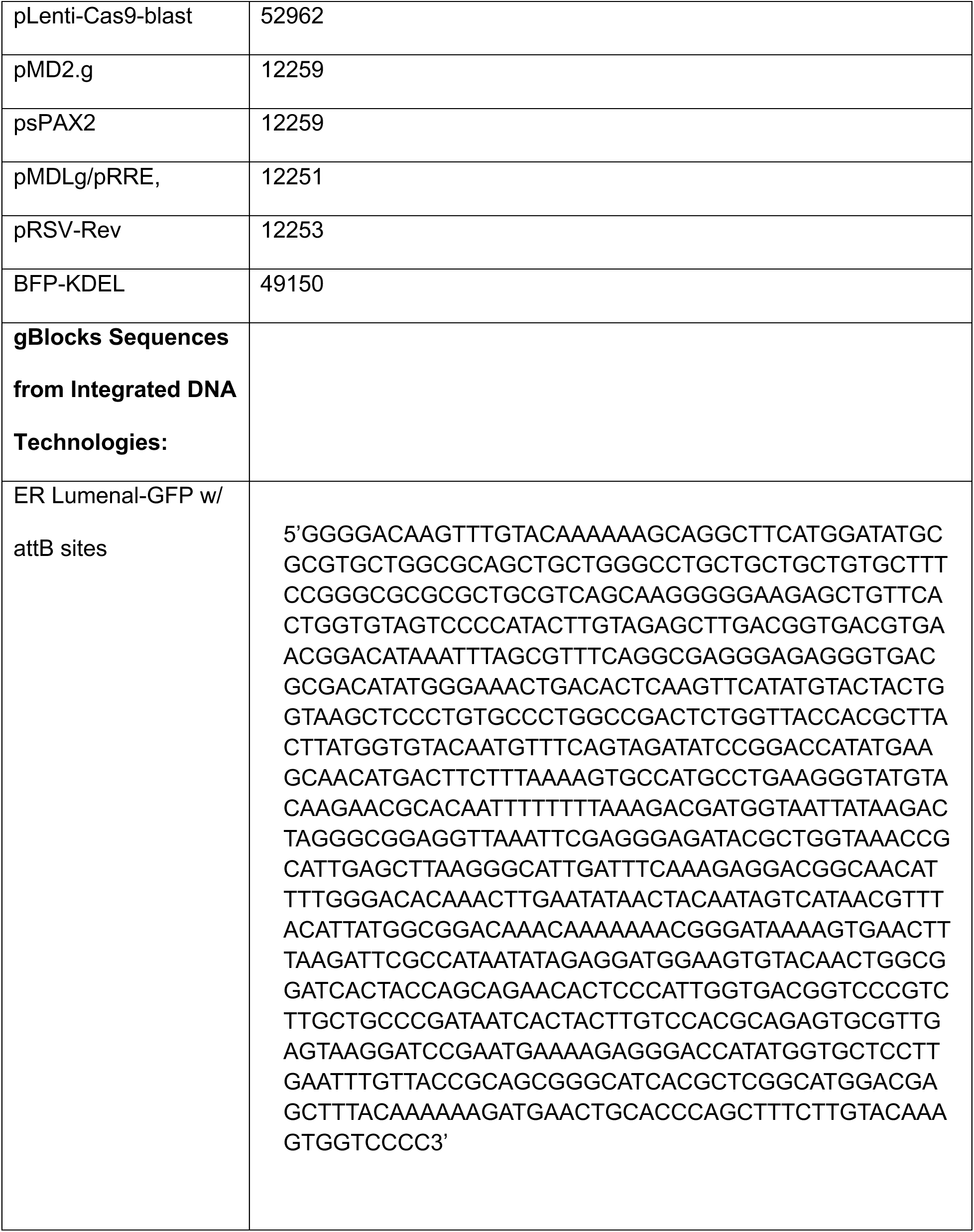
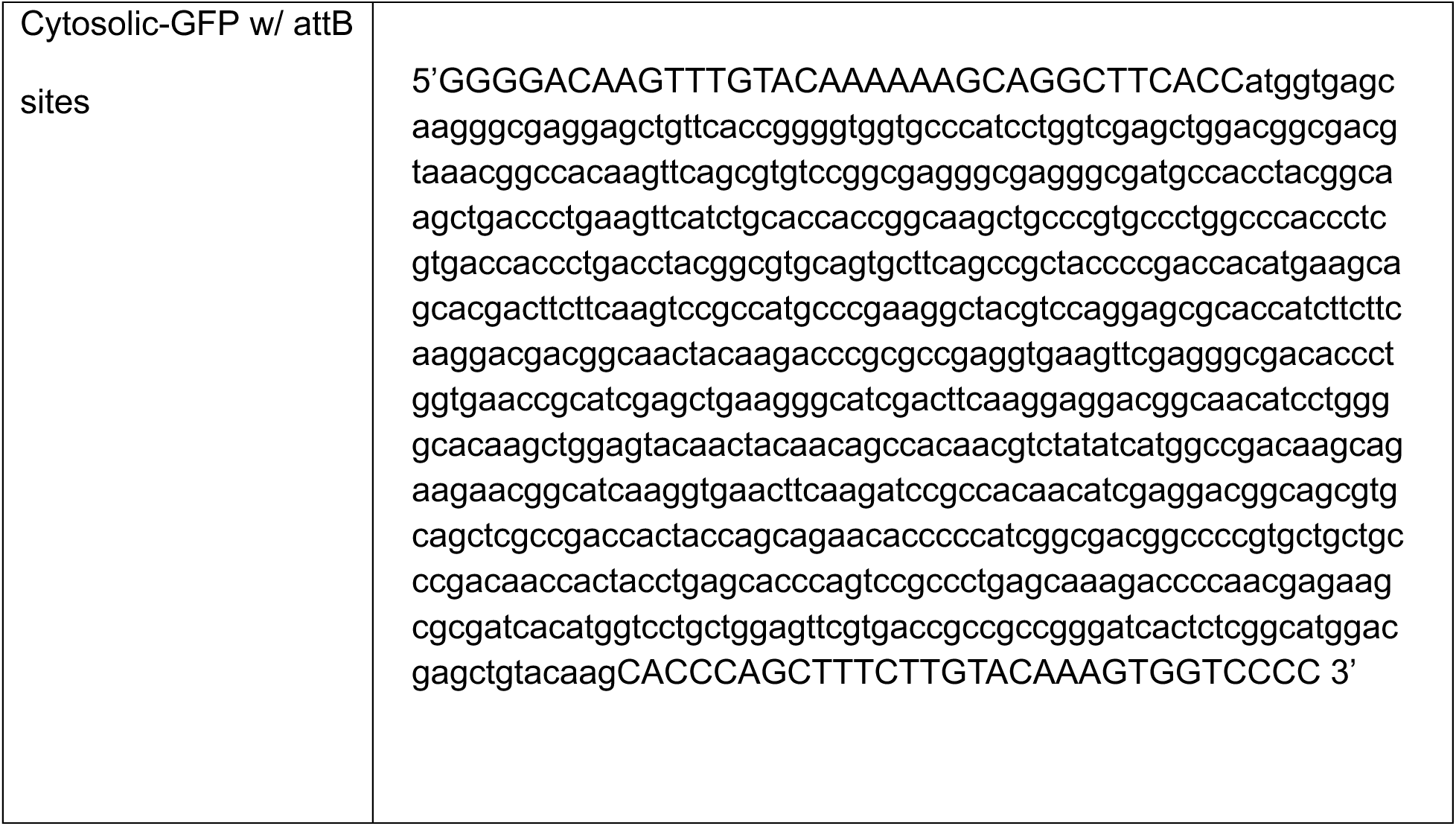
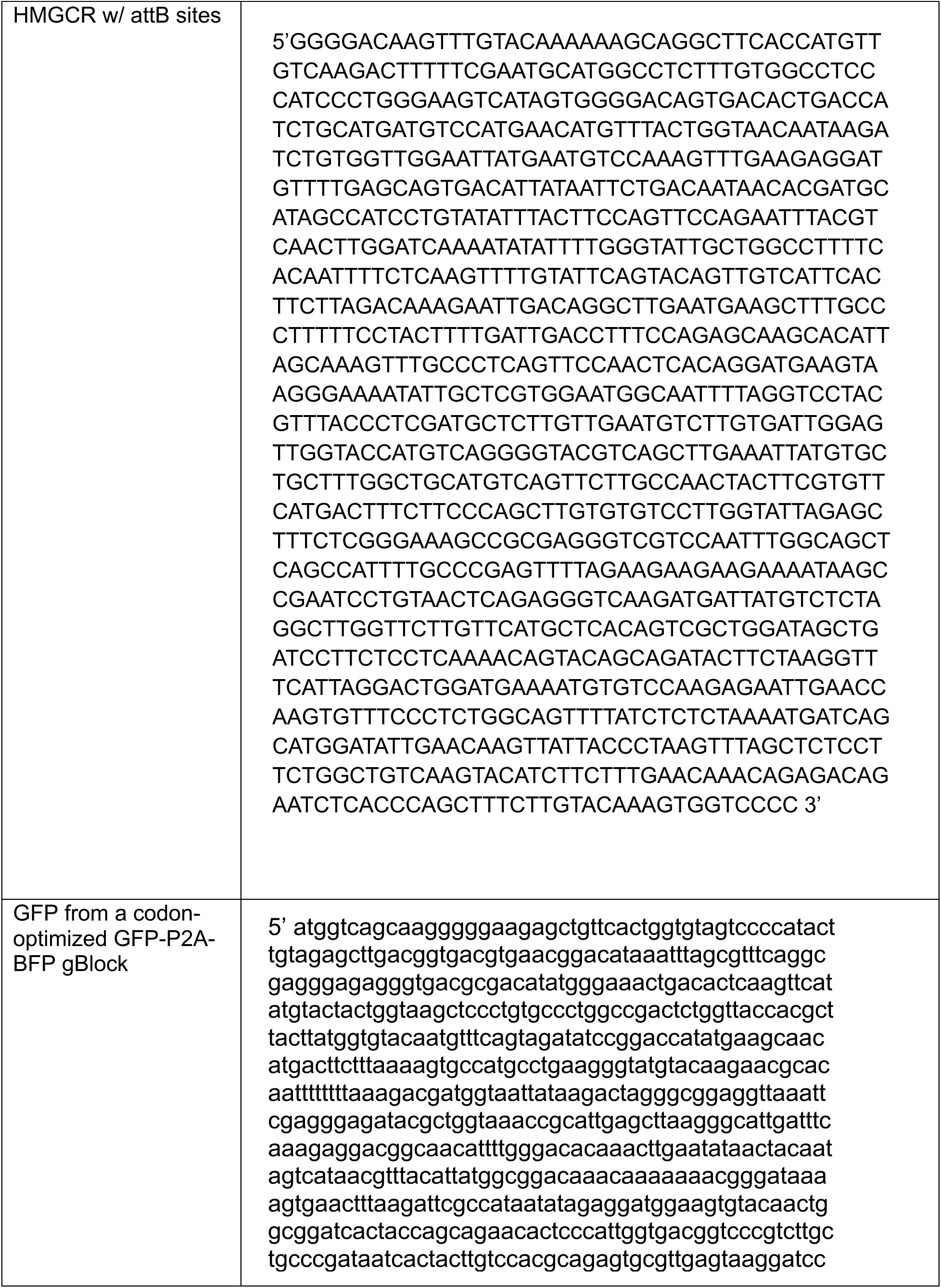
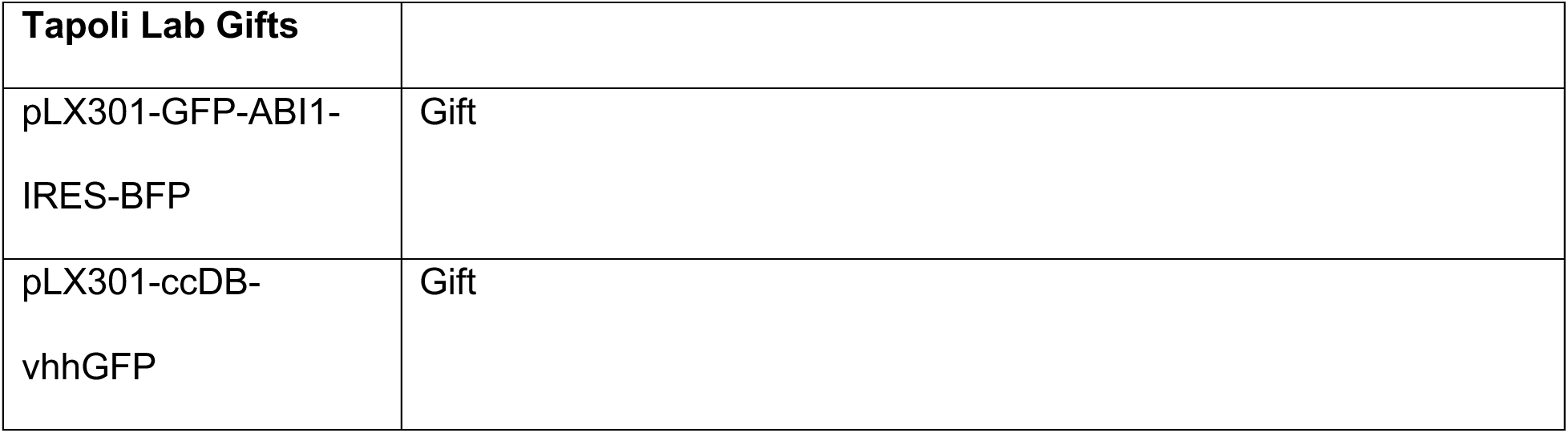

### Cell lines and culture conditions

HEK293T, U-2 OS, and Huh 7 cells were all obtained from UC Berkeley Cell Culture Facility. All cell lines were cultured in DMEM medium containing 4.5 g/L glucose, L-glutamine, without sodium pyruvate (Corning, no. 10-013-CMR). All media were supplemented with 10% fetal bovine serum (Gemini Bio Products) and all cells were maintained at 37°C and 5% CO_2_. Penicillin-streptomycin (Life Technologies, no. 15140122) was added to growth media for CRISPR-Cas9 screens and fluorescence-activated cell sorting. All cell lines were tested for mycoplasma.

### Generation of ER Membrane Protein-GFP-FKBP^F36V^ and Dual-tagged CANX-GFP-ABI1 with Effector-vhhGFP cell lines

Cytosolic GFP-FKBP^F36V^, ER Lumenal GFP-FKBP^F36V^, CANX-GFP-FKBP^F36V^, LDM-GFP-FKBP^F36V^, EPHX1-GFP-FKBP^F36V^, RTN4b-GFP-FKBP^F36V^, BCAP31-GFP-FKBP^F36V^, PKD2-GFP-FKBP^F36V^, and HMGCR-GFP-FKBP^F36V^ lines were generated by transduction with GFP-dTAG virus (as described in ‘Plasmids’) encoding the appropriate membrane protein, and second-generation lentiviral packaging plasmids. Cells were then selected in medium containing 2 ug/mL puromycin and analyzed by flow cytometry. CANX-GFP-FKBP^F36V^, LDM-GFP-FKBP^F36V^, EPHX1-GFP-FKBP^F36V^, and PKD2-GFP-FKBP^F36V^ U-2 OS cell lines were enriched via two rounds of FACS.

U-2 OS cells stably expressing GFP-tagged Calnexin were made via 2nd generation lentiviral transduction with a pLX301-CANX-GFP-ABI1-IRES-BFP plasmid, followed by selection with 6 ug/mL blasticidin, and FACS to enrich for GFP-signal. 2nd generation lentiviral transduction was performed to introduce each Effector-vhhGFP plasmid into the stable CANX-GFP-ABI1-IRES-BFP U2OS cell line resulting in 5 unique dual-tagged cell lines expressing both CANX-GFP and either VCP-vhhGFP, NPLOC4-vhhGFP, UFD1L-vhhGFP, CRBN-vhhGFP, or VHL-vhhGFP.

For the CRISPR-Cas9 genetic screens, U-2 OS Cytosolic GFP-FKBP^F36V^, CANX-GFP-FKBP^F36V^, and PKD2-GFP-FKBP^F36V^, lines stably expressing Cas9 were generated by transduction with pLenti-Cas9-blast (Addgene #52962) and third-generation lentiviral packaging plasmids. Cells were selected and maintained in medium containing 10 µg/mL or 4ug/mL blasticidin (Thermo Fisher Scientific #A1113903) for U-2 OS and HEK293T cell respectively. Cas9 activity was validated using the mCherry self-cutting system as previously described^40^. Briefly, cells were transfected with a plasmid expressing mCherry and a mCherry sgRNA, and mCherry fluorescence was assessed via flow cytometry after at least one week of growth.

Lentiviral particles for transduction were generated by co-transfection of lentiCRISPRv2 plasmids with second-generation lentiviral packaging plasmids (pMD2.g and psPAX2) or lentiCRISPRv3 plasmids with third-generation lentiviral packaging plasmids (pMDLg/pRRE, pRSV-Rev, and pMD2.G) into HEK293T cells using TransIT-LT1 transfection reagent (Mirus) according to manufacturer’s instructions. Lentiviral media was collected 72 h after transfection, passed through a 0.45 µm syringe filter, and used to infect U-2 OS and HEK293T cells in the presence of 8 µg/mL polybrene.

### Immunoblots

Cells were washed twice with phosphate-buffered saline (PBS), collected by scraping, spun down at 500g for 5 m at 4C. Cells were lysed in RIPA buffer supplemented with ETDA-free Pierce Protease and Phosphatase Inhibitor mini tablets (Thermo Scientific #A32961), sonicated for 15 s, recovered on ice for 2 m and, subjected to a second round of sonication and recovery. Cell lysate was then centrifuged for 10 min at 15,000*g* to remove any cell debris. Protein concentrations were determined using the bicinchoninic acid protein assay (Thermo Fisher Scientific), and equal amounts of protein by weight were combined with Laemmli buffer, boiled for 5 min at 95 °C (except when blotting for PKD2 and HMGCR), separated on 4–20% polyacrylamide gradient gels (Bio-Rad Laboratories) and transferred onto nitrocellulose membranes (Bio-Rad Laboratories). For ubiquitin blots, an overnight transfer was performed using wet transfer buffer with methanol onto PVDF membranes (Bio-Rab Laboratories). Membranes were blocked in PBS-containing or TBS-containing (for ubiquitin) 0.1% Tween 20 (PBST or TBST) containing 5% (w/v) bovine serum albumin (BSA) (Sigma-Aldrich) for 60 min. Membranes were incubated overnight at 4 °C in PBST or TBST containing 3% BSA and primary antibodies. After three washes with PBST or TBST, membranes were incubated at room temperature for 60 min in 3% BSA in PBST or TBST containing fluorescent secondary antibodies, followed by three more washes with PBST or TBST. Immunoblots were then imaged on an LI-COR imager (LI-COR Biosciences).

For the dual-tagged CANX-GFP-ABI1-IRES-BFP with effector-vhhGFP cell lines, effector-vhhGFP substrates were detected using the MonoRab HRP rabbit anti-camelid VHH antibody (GenScript, A01861). Immobilon Western Chemiluminescent HRP substrate (Millipore) was used to generate a chemiluminescence signal which was then detected with a ChemiDoc MP imaging system (BioRad).

The following blotting reagents and antibodies were used: anti-VCP (Novus, NB100-1558) anti-GFP (Sigma-Aldrich #SAB1305545), anti-HA (Sigma-Aldrich #H9658), anti-α-Tubulin (Cell Signaling #11H10), Anti-β-Actin (ACTB) Antibody (Sigma-Aldrich # A5441) UBXD8 (generated inhouse) anti-Ub (Enzo Life Sciences, no.BML-PW0930), IRDye800 conjugated secondary (LI-COR Biosciences, #926-32211, 1:20,000) and antimouse Alexa Fluor 680 conjugated secondary (Invitrogen, #A21058, 1:20,000). All primary antibodies were diluted in 1:1,000 unless stated otherwise.

### Flow cytometry

Fluorescence intensities were measured using an LSR Fortessa (BD Biosciences) at the UC Berkeley Flow Cytometry Core Facility. U-2 OS or HEK293T cells expressing ER Membrane Protein–GFP–FKBPF36V or dual-tagged CANX–GFP–ABI1 with effector-vhhGFP constructs were seeded in 12-well plates in DMEM supplemented with 10% FBS.

Cells were dissociated from plates using TrypLE Express (Gibco) and resuspended in DMEM containing 10% FBS. Cells were pelleted by centrifugation at 500 × g for 5 min, resuspended in DPBS, and placed on ice. Samples were vortexed and analyzed at 1,000 events/s for a total of 10,000 events. Live cells were gated using SSC-A vs. FSC-A, and single cells were gated using FSC-H vs. FSC-A. GFP signal was detected using the 488 nm laser, and mean fluorescence intensity (MFI) was calculated in FlowJo v10. To account for background fluorescence, U-2 OS or HEK293T wild-type cells lacking the ER Membrane Protein–GFP–FKBPF36V reporter were used as controls. The average MFI from control cells was subtracted from the reporter MFI at all time points and doses to calculate percent GFP remaining.

### Dose-Response

U-2 OS and HEK293T ER-TM Protein–GFP–FKBP^F36V^ cells were seeded in DMEM supplemented with 10% FBS, and Huh7 cells were seeded in DMEM containing 10% lipoprotein-deficient FBS, in p60 dishes or 12-well plates for analysis by immunoblot or flow cytometry, respectively. 24 h later, cells were treated with or without small-molecule (dTAGv-1 (#), dTAG-13 (Sigma-Aldrich #SML2601), atorvastatin, or 11j/P22A) at the indicated concentrations for 24 h. Cell harvest and sample preparation for immunoblotting and flow cytometry were performed as described above. For immunoblot analysis, total protein lysates were prepared as described above, and 35 µg (U-2 OS), 10 µg (HEK293T), or 30 µg (Huh7) was loaded per lane.

### Emetine-Chase Assays

U-2 OS ER Membrane Protein–GFP–FKBP^F36V^ cells were seeded in 12-well plates and treated with 75 µM emetine (Sigma-Aldrich #E2375-50MG), 100 nM dTAGv-1 (for cytosolic GFP), 500 nM dTAGv-1 (for all other ER transmembrane proteins), or combinations thereof at indicated times. Cell harvest and sample preparation flow cytometry were performed as described above.

### VCP Inhibitor Experiments

U-2 OS ER Membrane Protein–GFP–FKBP^F36V^ cells were seeded in p60 dishes or 12-well plates for immunoblot or flow cytometry, respectively. Cells were pretreated for 30 min with 5 µM CB-5083 (Cayman #C814B92), 25 µM NMS-873 (Fisher Scientific #501015294), or 500 nM MLN4924 (#CT-M4924), followed by co-treatment with the same inhibitor and 500 nM dTAGv-1 or dTAG-13, or degrader alone at the indicated time points. DMSO served as the vehicle control.

Huh7 cells were seeded in DMEM supplemented with 10% lipoprotein-deficient FBS in p60 dishes for immunoblot analysis 24 h prior to treatment. Cells were pretreated for 30 min with 5 µM CB-5083 or 500 nM MLN4924, followed by co-treatment with the same inhibitor and 0.3 µM 11j/P22A, or degrader alone for 24 h. DMSO and atorvastatin-only treatments were included as controls. Cells were harvested, and lysates prepared for immunoblotting or flow cytometry as described above. For immunoblot analysis, 35 µg (U-2 OS) and 30 µg (Huh 7) total protein was loaded per lane.

### Proteasome and Lysosome Inhibitor Assays

U-2 OS ER Membrane Protein–GFP–FKBP^F36V^ cells were seeded in 12-well plates for analysis flow cytometry. For proteasome and lysosome inhibitor assays, cells were pretreated for 3 h with either 10 µM MG132 (Selleck #S2619) or 250 nM Bafilomycin A1 (Sigma-Aldrich #B1793), followed by co-treatment with the same inhibitor and dTAGv-1, or dTAGv-1 alone, at indicated times and doses. Cell harvest and sample preparation for flow cytometry were performed as described above.

### Time Course

Huh7 cells were seeded in DMEM containing 10% lipoprotein-deficient FBS, in p60 dishes, and 24 h later, treated with 0.3uM of 11j/P22A or atorvastatin (Spectrum Chemical & Laboratories Products #TCI-A2476) at the indicated time points. Cell harvest and sample preparation for immunoblotting were performed as described above with 30ug of total protein lysate loaded per lane.

### Fluorescence microscopy

For super-resolution microscopy of live cells, ER Membrane Protein-GFP-FKBP^F36V^ cells were seeding into 12-well plate, and 24 h later transiently transfected using X-tremeGENE (Millipore Sigma #06366546001) with BFP-KDEL (Addgene #49150) for imaging the ER according to manufacturing recommendations. The next day, cells were seeded into 4-well or 8-well Nunc™ Lab-Tek™ II Chambered Coverglass (Borosilicate Glass 1.5; Thermo Fisher Scientific, 155360). If treated, the indicated dTAGv-1 or DMSO dose was added 24 h before the start of imaging. Prior to imaging, cells were washed 2x with DPBS and imaged in fresh phenol red-free medium supplemented with 10% FBS. Live cells were imaged using a Zeiss Axio Observer 7 fitted with a 63X oil objective using DAPI, and GFP filters. Cells were imaged at 37 °C with 5% CO_2_. Z-stacks of 0.2-μm thickness were acquired.

### Generation of custom UBALER sgRNA library

The UBALER library is a proportionally equivalent combination of the UBAL library and a custom ERADplus library. The previously published UBAL sgRNA library includes 20,710 elements: 18,710 sgRNAs targeting 1,871 genes (∼10 sgRNAs per gene) and 2,000 negative control sgRNAs^41^. The custom ERADplus sgRNA library contains 5,564 elements with 3,564 sgRNAs targeting 389 genes (∼10 sgRNAs per gene) and 2,000 negative control sgRNAs. Guide sequences were from the Bassik Human CRISPR Knockout Library, and the library construction protocol was previously described^19,42^. Therefore, the UBALER library consists of 26,274 elements: 22,274 sgRNAs targeting 2,260 genes (∼10 sgRNAs per gene) and 4,000 negative control sgRNAs. The UBALER library targets genes involved in ubiquitin conjugation, deubiquitination, ubiquitin-like conjugation, proteasome, autophagy and lysosome pathways, ERAD and glycosylation machinery.

### UBALER U-2 OS Cytosolic GFP-FKBP^F36V^, CANX-GFP-FKBP^F36V^, PKD2-GFP-FKBP^F36V^ CRISPR-Cas9 screens

To generate lentiviral particles, the UBALER library was co-transfected with third-generation lentiviral packaging plasmids into HEK293T cells. Media containing lentivirus was collected 72 h after transfection, and filtered. U-2 OS Cytosolic GFP-FKBP^F36V^, CANX-GFP-FKBP^F36V^, and PKD2-GFP-FKBP^F36V^ stably expressing Cas9 were transduced with lentiviral packaged UBALER library in 8 µg/mL polybrene to achieve a 20%–50% mCherry-positive population, ensuring at least 200× coverage. Viral media was removed after 24 hours, and cells were expanded at 1000x coverage.

On the day of sorting, cells were treated with 100 nM dTAGv-1 for 15 h and 11 min (Cytosolic GFP-FKBP^F36V^), or 500 nM dTAGv-1 for 1 h and 37 min (CANX-GFP-FKBP^F36V^) and 50 mins (PKD2-GFP-FKBP^F36V^), dissociated using TrypLE™ Express Enzyme (Gibco #12605010), collected by centrifugation (500 × g for 5 minutes), washed with 1x DPBS (Gibco #1419-144), and resuspended in phenol red-free media (HyClone #16777-406) supplemented with 3% FBS and 1% fatty acid-free BSA. Cells were passed through 70-μm cell strainers (Falcon #352350) and kept on ice until FACS.

Cells were sorted using a BD FACS Aria Fusion Cell Sorter (BD Biosciences) at the UC Berkeley Flow Cytometry Core Facility. The top 5% and bottom 75% GFP+ cells were isolated using the following gating hierarchy: SSC-A v FSC-A polygonal gate for live cells, FSC-H v FSC-A polygonal gate for singlets, a histogram gate for mCherry+ cells using the 561 nm yellow-green laser, and a histogram gate for the top 5% and bottom 75% GFP+ cells using the 488 nm blue laser. Samples were kept at 4°C and sorting was performed using a 85 μm nozzle at a flow rate of 8-10,000 events/s, with four-way purity. Approximately 1.32 × 10⁶ top 5% and 20 × 10⁶ bottom 75% GFP+ cells were collected to ensure 1,000× coverage of the ∼26,000-element UBALER library.

Genomic DNA was extracted using the QIAamp DNA Blood Midi Kit (Qiagen) as per the manufacturer’s instructions. Guide sequence libraries were prepared from genomic DNA by two rounds of PCR using Herculase II Fusion DNA Polymerase (Agilent #600679)^40^. First, sgRNAs were amplified using the following reaction mix for a 100 μL reaction: 10 μg genomic DNA, 5x Herculase buffer (20 μL), 100 μM primers oMCB_1562 (1 μL) and oMCB_1563 (1 μL), 100 mM dNTPs (1 μL), 2 μL Herculase II Fusion DNA Polymerase, and nuclease-free water. PCR conditions were: 1x 98°C (2 min); 18x 98°C (30 sec), 59.1°C (30 sec), 72°C (45 sec); 1x 72°C (3 min). Next, amplicons were indexed using Illumina TruSeq LT adapters. The following reaction mix was used for indexing: 10 μL PCR1 product, 5x Herculase buffer (20 μL), 100 μM primers oMCB_1439 (0.8 μL) and barcoded oMCB_1440 (0.8 μL), 100 mM dNTPs (2 μL), 2 μL Herculase II Fusion DNA Polymerase, and nuclease-free water. PCR conditions were: 1x 98°C (2 min); 20x 98°C (30 sec), 59.1°C (30 sec), 72°C (45 sec); 1x 72°C (3 min).

PCR products were separated on a 2% TBE-agarose gel, purified using the QIAquick Gel Extraction Kit (Qiagen #28704), and assessed for quality using a Fragment Analyzer (Agilent). Amplicons from each cell line (GFPhigh and GFPlow) were pooled based on concentrations (as determined by Qubit Fluorometric Quantification) and the number of elements in the UBALER library. The sgRNA sequences were analyzed by deep sequencing on an Illumina NextSeq instrument at the UC Berkeley QB3 Functional Genomics Laboratory, using the standard Illumina indexing primer and custom sequencing primer oMCB_1672^40^.

Sequence reads were aligned to the sgRNA reference library using Bowtie 2 software. For each gene, the gene effect and score (representing the likely maximum effect size and corresponding score) were calculated using the Cas9 high-throughput Maximum Likelihood Estimator (casTLE) statistical framework as previously described^19,42^.

Primer sequences:

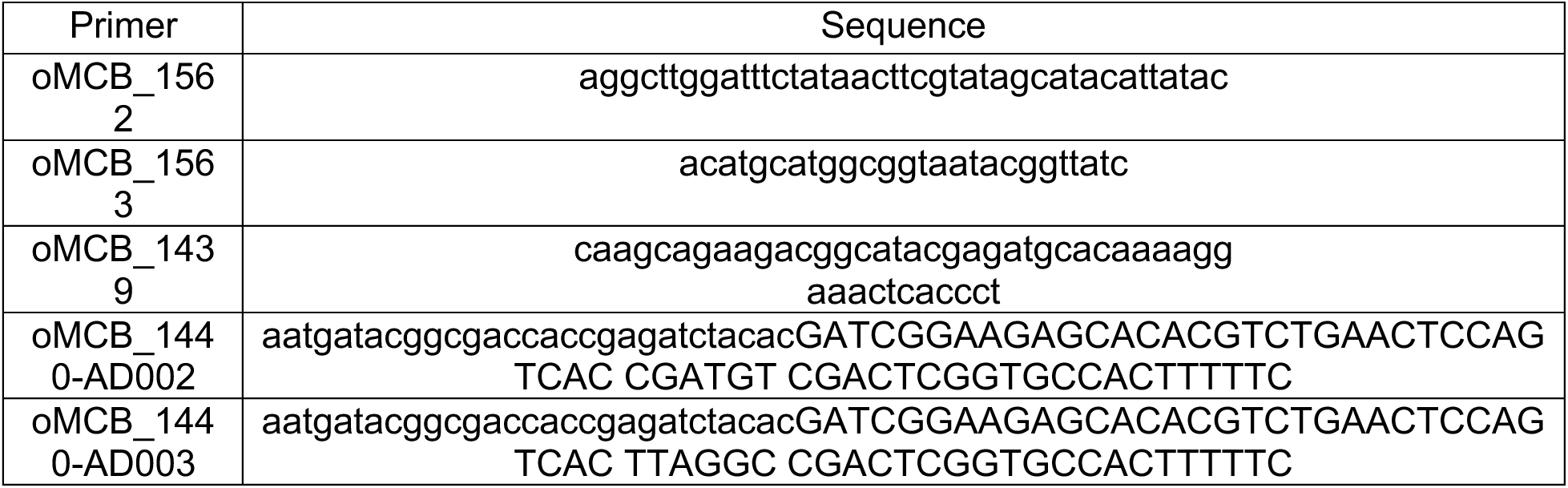
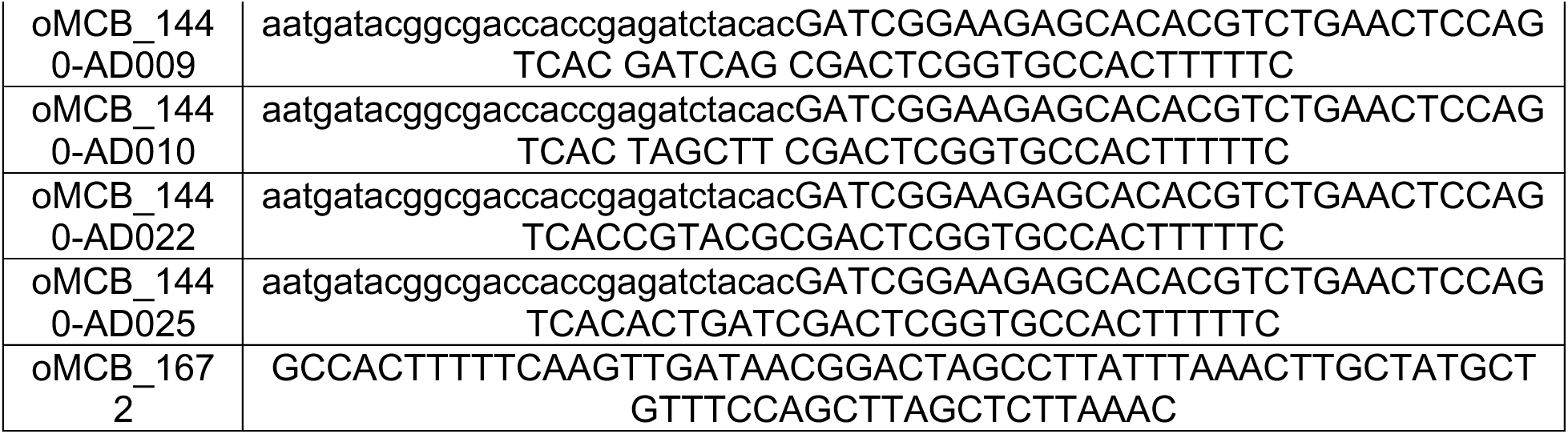

### Immunoprecipitations

U-2 OS CANX-GFP-FKBP^F36V^ or HMGCR-GFP-FKBP^F36V^ cells were seeded into 15-cm^2^ plates. The following day, CANX-GFP-FKBP^F36V^ cells were pre-treated for 30 min with either 5 mM CB-5083 and 500 nM MLN4924, 5 mM CB-5083 alone, or DMSO, followed by a 90 min treatment with 500 nM dTAGv-1 in the continued presence of the pre-treatment conditions. Cells were washed twice with PBS, collected by scraping, spun down at 500g for 5 min at 4°C and frozen at -80 C. Cell pellets were thawed on ice and resuspended in 600 µL lysis buffer (50 mM Tris-HCl pH 7.5, 150 mM NaCl, 0.5 mM EDTA, 1.0% Digitonin (Neta Scientific Inc #SIAL-300410), ETDA-free Pierce Protease and Phosphatase Inhibitor mini tablets (Thermo Scientific #A32961), 10 mM N-ethylmaleimide (NEM; Thermo Fisher Scientific # E1271-1G), and 10 uM MG132 (Selleck #S2619)). Lysates were sonicated for 15 sec, recovered on ice for 2 min and, subjected to a second round of sonication and recovery. Cell lysate was then centrifuged for 10 min at 15,000*g* to remove any cell debris. Protein concentrations were determined using the bicinchoninic acid protein assay (Thermo Fisher Scientific). 1 mg of lystates were diluted with dilution buffer (50mM Tris-HCl pH 7.5, 50 mM sodium chloride, 0.5 mM EDTA (adjust the pH at 4°C), and protease inhibitor) and incubated with 10 µL of equilibrated ChromoTek GFP-Trap Magnetic Agarose (Proteintech, no. gtma) for 1 h at 4°C with end-over-end rotation. Beads were washed 3 times with washing buffer (50 mM Tris-HCl pH 7.5, 50 mM sodium chloride, 0.5% IGEPAL) and eluted in 10 µL of 2X Laemmli buffer at 65°C for 15 min. Eluates were run on 4–20% polyacrylamide gradient gels (Bio-Rad), transferred to nitrocellulose or PVDF membranes, and immunoblotted as described in the ‘immunoblotting’ methods.

For DUB digestions, beads were washed twice with 500 mL DUB buffer (50 mM Tris pH 7.5, 50 mM sodium chloride, 2 mM DTT), resuspended in DUB buffer, and incubated with 150 nM of Recombinant Human His6-USP2 Catalytic Domain Protein Catalytic Domain (Bio-Techne #E-506) at 30°C for 1 hr with gently shaking. Supernatant was removed and resuspend in 10uL of 2X Laemmli buffer at 65 °C for 15 min for elution.

For proteomics, beads were washed three times with 500 µL of 50 mM ammonium bicarbonate (ABC, pH 8.0), resuspended in 50 µL of 50 mM ABC containing 0.1% RapiGest (Waters #186008090). Beads were vortexed, trypsin was added at a 1:100v/v ratio, and allowed to incubate overnight with shaking at 37°C. Samples were quenched with 5% TFA, desalted using a C18 desalting column (Waters, Sep pak 2cc 50mg), and dried using a speed-vac.

### Proteomics

Peptides were resuspended in 1% formic acid and separated on an Easy nLC 100 UHPLC equipped with a 15 cm nano-liquid chromatography column. Using a flow rate of 300 nL/min, the linear gradient was 5% to 35% over B for 90 min, 35% to 95% over B for 5 min and 95% hold over B for 15 min (solvent A: 0.1% formic acid in water, solvent B: 0.1% formic acid in ACN). Precursor ions were acquired (m/z 350-2000) at 120K resolving power with an isolation window of 10 ppm. High-energy collisional dissociation (HCD) with 27% collision energy was collected using the ion trap with an automatic gain control (AGC) target of 1e4 and an isolation window of 2 m/z. Peptides were identified and relative abundances were determined using Proteome Discoverer 2.4. Results are represented as average ± s.d. of duplicates.

### Differential Fractionation

15-cm^2^ plates were seeded, treated, and collected for each condition. Cultured cells were collected, washed with ice-cold phosphate-buffered saline (PBS), and incubated in hypotonic lysis medium (HLM: 20 mM Tris-HCl, pH 7.4, 1 mM EDTA) supplemented with 10 mM NEM (Thermo Fisher Scientific # E1271-1G) on ice for 10 min. 23G needle was used to lyse the cells by pushing through the needle 8 times. Samples were centrifuged (500 × g, 5 min, two times) to remove unbroken cells. The remaining supernatant was then centrifuged (20,000 × g, 30 min at 4°C) to separate heavy membrane and cytosolic fractions. The resulting pellet (membrane) was then reconstituted to its corresponding cytosolic fraction volume using HLM buffer. For immunoblotting, BME was then added to achieve a final detergent concentration of 1% and equal volumes were analyzed.

### Dual-tagged CANX-GFP-ABI1-IRES-BFP with effector-vhhGFP cell lines degradation assessment

For dual-tagged CANX-GFP-ABI1-IRES-BFP with effector-vhhGFP cell lines, cells were seeded, harvested, and prepared for flow cytometry as described above. Live, single cells were gated as previously described, with an additional SSC-W vs. SSC-H gate applied. GFP and BFP fluorescence were detected using 468 nm and 405 nm lasers, respectively. In the parental CANX-GFP-ABI1-IRES-BFP line, a low GFP population gate was defined on a BFP (x-axis) vs. GFP (y-axis) plot such that 1-5% of cells fell within the gate; this gate was then applied to all effector-vhhGFP lines **(Fig. S5C)**. MFI for GFP and BFP were calculated in FlowJo v10, and the GFP/BFP ratio determined. Fold change was calculated by dividing the parental GFP/BFP ratio by that of the effector-vhhGFP cell line. Graphs were generated using GraphPad Prism (v8–10) with default settings.

### Synthesis of 11j/P22A

11j/P22A **3 (Supplementary Fig. S8)** (3R,5R)-7-(3-(3-(3-(1-(14-((2-(2,6-Dioxopiperidin-3-yl)-1,3-dioxoisoindolin-4-yl)amino)-3,6,9,12-tetraoxatetradecyl)-1H-1,2,3-triazol-4-yl)propoxy)phenyl)-2-(4-fluorophenyl)-5-isopropyl-4-(phenylcarbamoyl)-1H-pyrrol-1-yl)-3,5-dihydroxyheptanoic acid was synthesized based on previously published procedure^12^. To a vial containing a stir bar were added 2 mL of THF, pomalidomide-4PEG-N3 1 (**Supplementary Fig. S8**) (12.1 mg, 23.3 μmol), and alkyne 2 (Fig 6SA) (12.5 mg, 19.4 μmol). To the same vial was then added 1 mL aqueous solution of Cu2SO4•5H2O (1.0 mg, 3.9 μmol) and sodium ascorbate (0.77 mg, 3.9 μmol). The mixture was stirred at room temperature overnight. The next day, THF in the crude mixture was removed under reduced pressure and then resuspended in 1 mL of DMSO. After filtration, crude mixture was purified by preparative HPLC with a 50-min gradient of 20% to 70% MeCN in water with 0.1% of formic acid additive and 9 mL/min constant flow rate. Product 3 (Fig 6SA) was detected by UV absorption at 280 nm. 3 (Fig 6SA) was isolated as a white solid (7.5 mg, 19.4 μmol, 33%, RT = 40 mins). H-NMR of the isolated compound 3 **(Supplementary Fig. S8**) matches previously reported data^12^

HRMS: calc. C61H72FN8O14 [M + H] +, 1159.5147; found, 1151.5091.

Instrumentation:

Prep-HPLC: Thermo Dionex UltiMate 3000 UHPLC+ Focused.

Column: Luna® 10 μm C18(2) 100 Å LC Column 250 x 21.2 mm (P/No. 00G-4253-P0-AX)

### RNA Isolation and Quantitative Real-Time PCR (TaqMan)

Total RNA was extracted from cells using the RNeasy Mini Kit (Qiagen) according to the manufacturer’s instructions. 2.5ug of total RNA was used for cDNA synthesis. cDNA was generated using the Maxima First Strand cDNA Synthesis Kit (Thermo Fisher Scientific, K1641) following the manufacturer’s protocol, which includes a DNase treatment step to remove residual genomic DNA. Reverse transcription reactions were carried out in a 20 µL volume with random hexamer and oligo(dT) primers under the following conditions: 25 °C for 10 min, 50 °C for 30 min, and 85 °C for 5 min to terminate the reaction.

Quantitative real-time PCR was performed using TaqMan Gene Expression Assays (Applied Biosystems) on a Applied Biosystems™ StepOnePlus™ Real-Time PCR System (Thermo Fisher Scientific #4376600) (Each 20 µL reaction contained 10 µL of TaqMan Universal PCR Master Mix II (2X), 1 µL of the gene-specific TaqMan probe/primer mixture (20X), 2 µL of cDNA (equivalent to ∼100 ng input RNA), and 7 µL of nuclease-free water. All samples were run in triplicate along with no-template controls and no-RT controls to monitor contamination and genomic DNA carry-over. Thermal cycling conditions were: 50 °C for 2 min (for UNG activation, if applicable), 95 °C for 10 min to activate DNA polymerase, followed by 40 cycles of 95 °C for 15 s (denaturation) and 60 °C for 1 min (annealing/extension).

Gene expression levels were normalized to the endogenous control gene GAPDH, and relative quantification was performed using the ΔΔCₜ method.

### Statistical analysis and reproducibility

All figures, including immunoblots, dose-response curves, protein turnover assays and panels are representative of three biological replicates unless stated otherwise. For comparison across multiple experimental groups, p-values were calculated using one-way ANOVA and adjusted using the Bonferroni correction for multiple comparisons (****P < 0.0001, ***P < 0.001, **P < 0.002, *P < 0.033).

---

**Supplementary Fig. S1.**
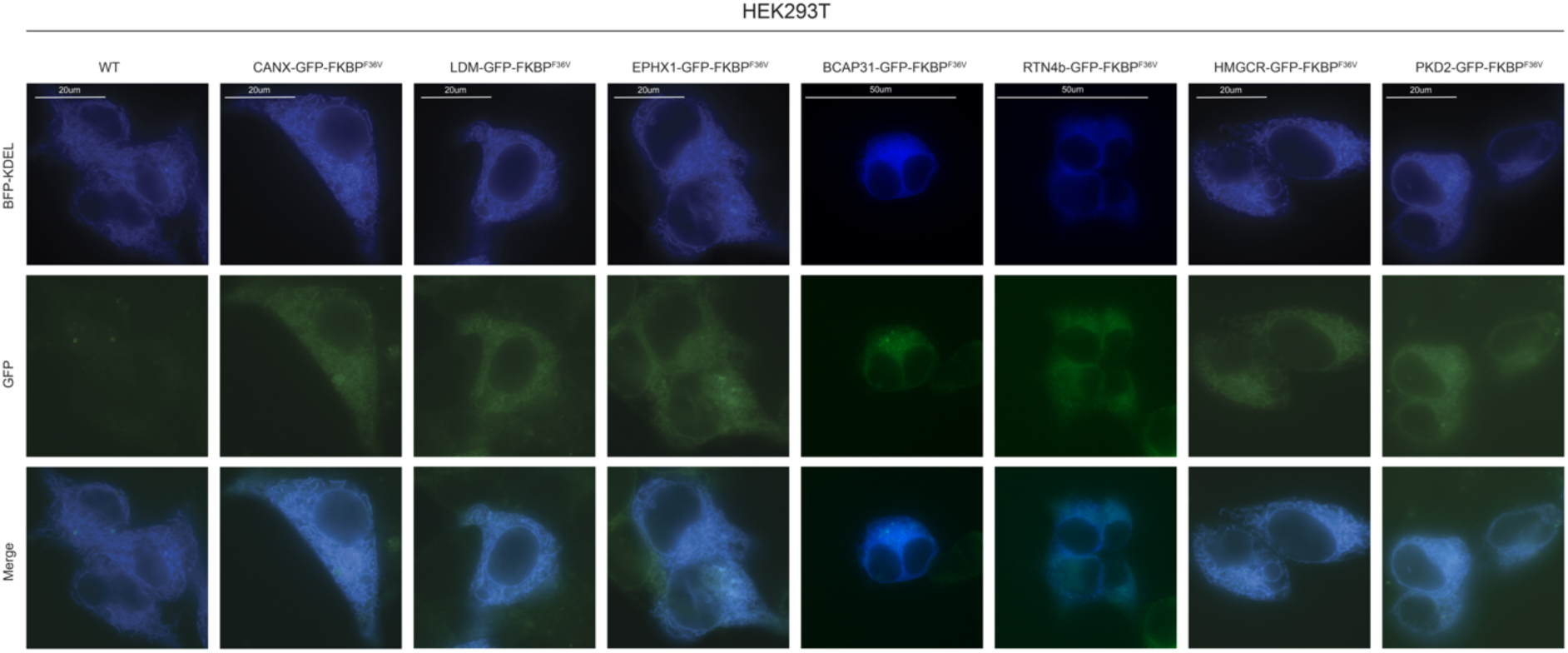
HEK293T ER protein reporter cell lines co-localized to the ER. Fluorescence microscopy of GFP-tagged ER-TM reporters (green) co-localized with the ER marker BFP-KDEL (blue) in HEK293T cells. Scale bar, 20 and 50 μm.

**Supplementary Fig. S2.**
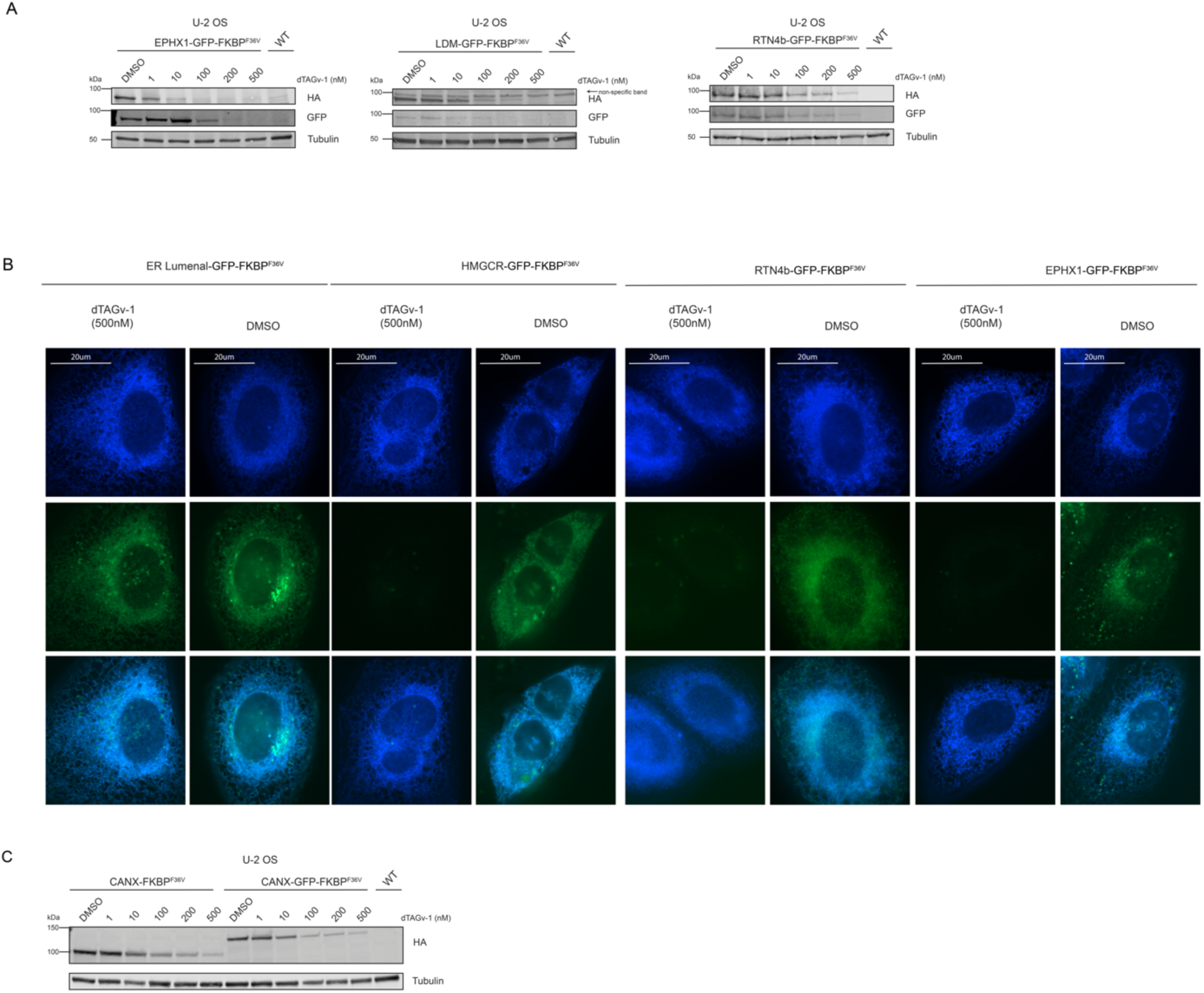
Induced degradation of ER proteins in U-2 OS cells. **(A)** The indicated fluorescent protein reporter cell lines were treated with the specified concentrations of dTAGv-1 or dTAG-13 for 24 h and immunoblotted for HA, GFP, and α-tubulin (loading control). **(B)** Reporter cell lines transiently expressing the ER marker BFP-KDEL (blue) were treated with the indicated concentrations of dTAGv-1for 24 h and imaged by fluorescence microscopy. GFP fluorescence (green) indicates the expressed proteins. Scale bar, 20 µm. **(C)** 24 hr dose-response analyzed by immunoblotting for HA or α-tubulin in specified cell lines (n=1).

**Supplementary Fig. S3.**
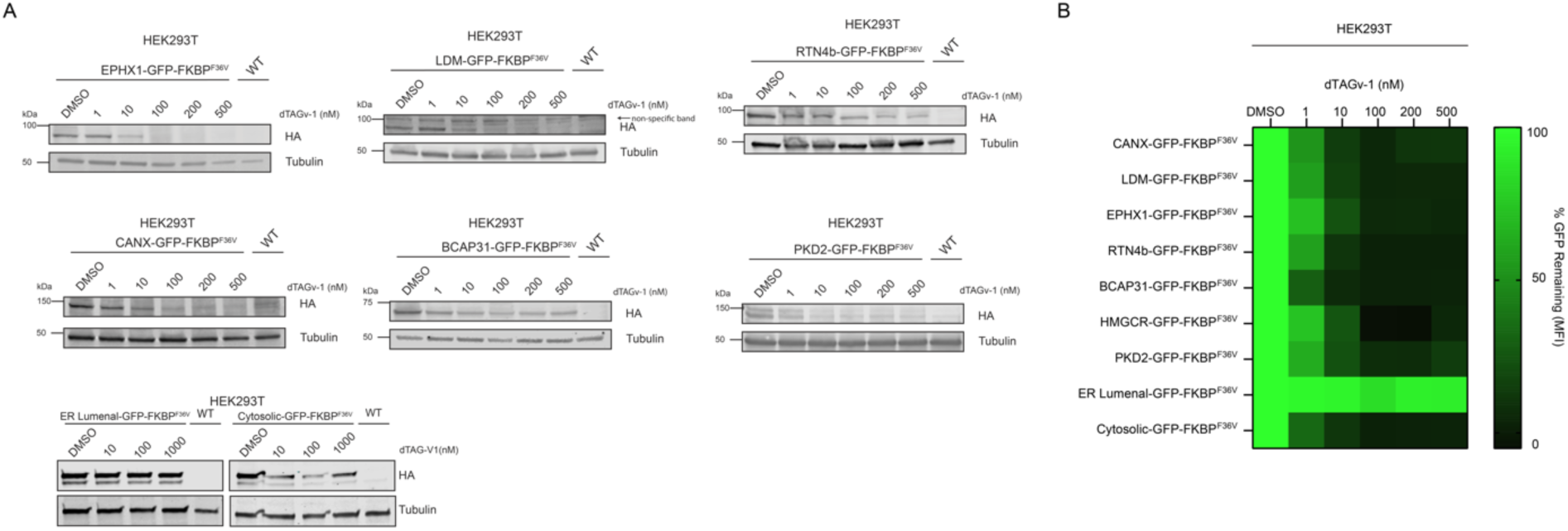
Induced degradation of ER proteins in HEK293T cells. **(A)** The indicated fluorescent reporter cell lines were treated with the specified concentrations of dTAGv-1 for 24 hr and immunoblotted for HA and α-tubulin (loading control). **(B)** GFP fluorescence was measure using flow cytometry following treatment with increasing doses of dTAGv-1 and the % GFP remaining visualized as a heatmap.

**Supplementary Fig. S4.**
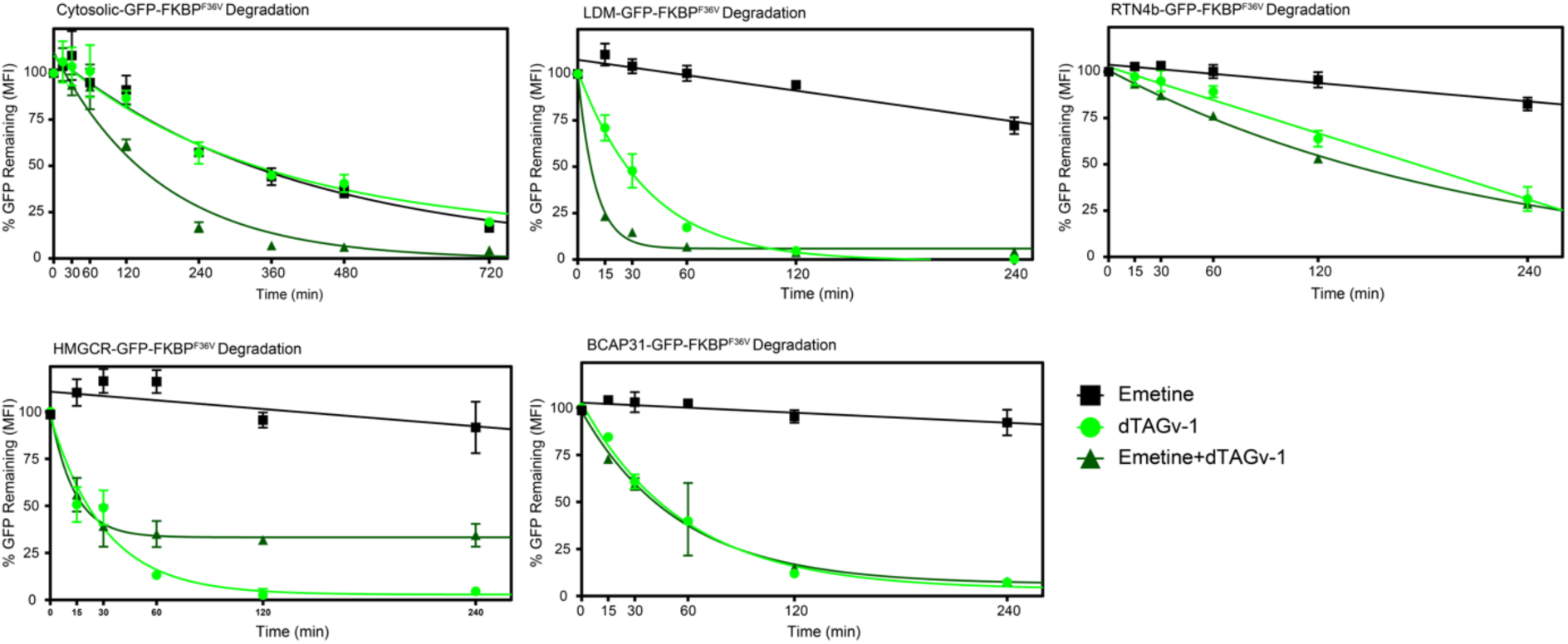
Analyses of substrate turnover kinetics. Flow cytometry was used to analyze U-2 OS reporter cell lines treated with 75 µM emetine, 100 nM dTAGv-1 (for cytosolic GFP-FKBP^F36V^), 500 nM dTAGv-1 (for all other ER-TM proteins), or the indicated combinations. GFP fluorescence was measured by flow cytometry.

**Supplementary Fig. S5.**
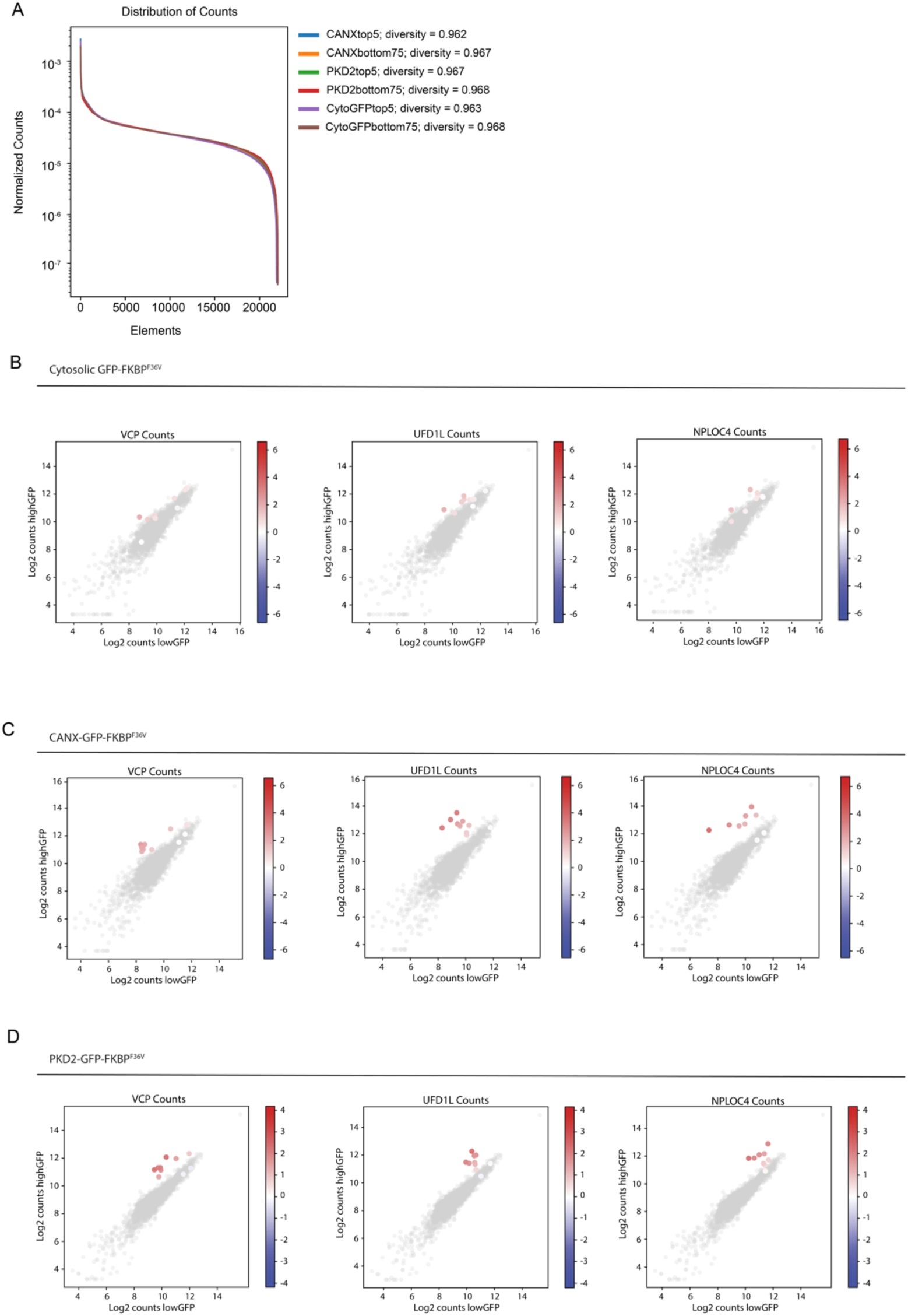
Genetic screens for mechanism of cytosolic E3 ligase-induced degradation of ER-TM proteins. **(A)** Distribution of counts across all sgRNA elements cloned into the custom UBALER sgRNA library. **(B-D)** Cloud plots indicating count numbers corresponding to *VCP, UFD1L,* and *NPLOC4* (color scale) and control (gray scale) sgRNAs for each screen.

**Supplementary Fig. S6.**
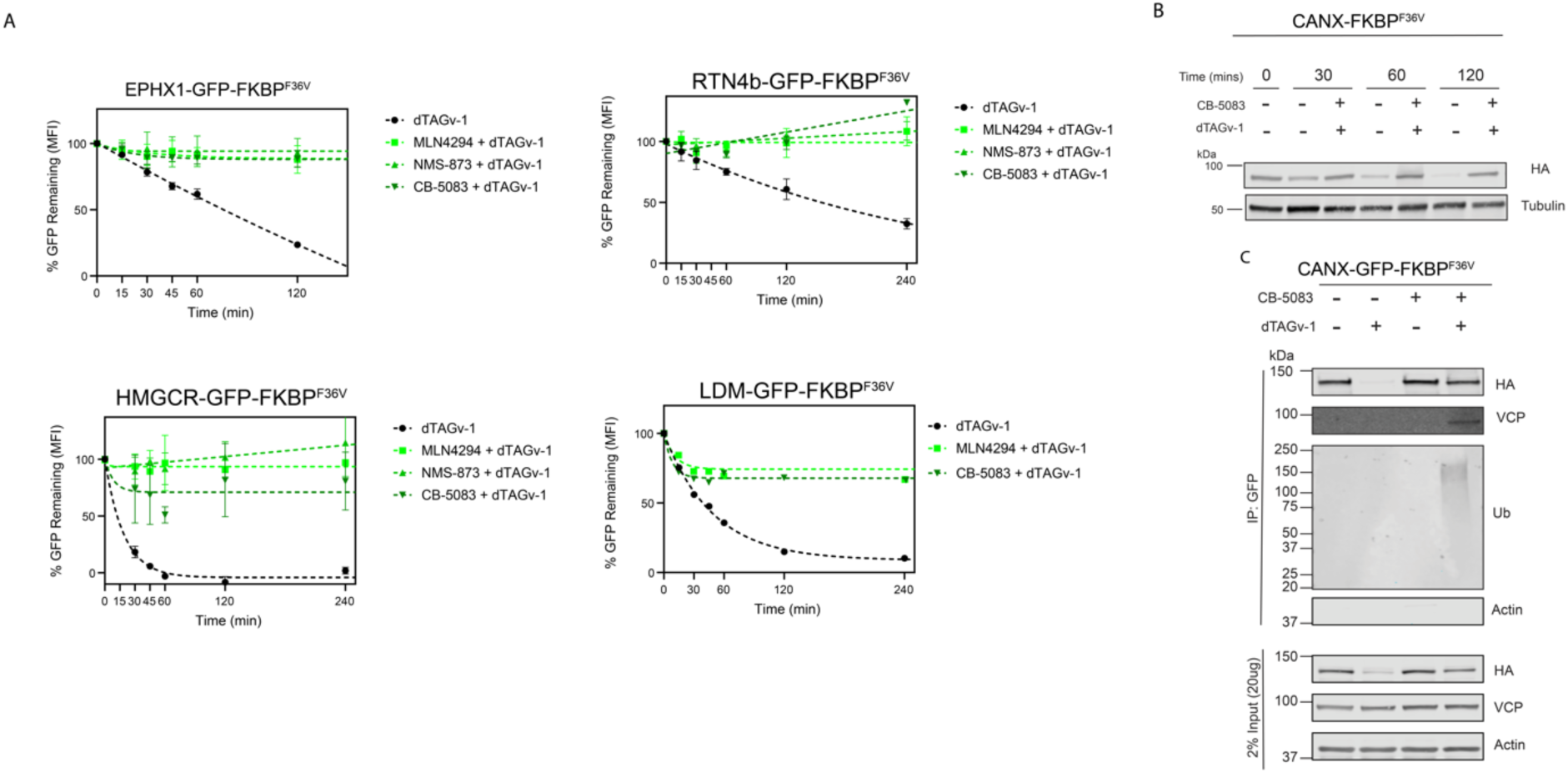
ER-TM protein degradation by recruitment of cytosolic E3 ligase requires ubiquitin and VCP activity. **(A)** Specified U-2 OS fluorescent ER-TM protein reporter cell lines along with CANX-FKBP^F36V^ were pre-treated for 30 min with 5 µM CB-5083, 25 µM NMS-873, or 500 nM MLN4924, followed by co-treatment with the same inhibitor and 500 nM dTAGv-1 or dTAG-13 for the indicated times unless otherwise noted; DMSO served as the vehicle control. **(A)** Kinetics and quantification of GFP fluorescence decay in the indicated reporter cell lines. **(B)** Immunoblot of CANX-FKBP^F36V^ cells probed for HA-tagged protein and α-tubulin. **(C)** GFP immunoprecipitation (IP) followed by immunoblotting for HA, VCP, ubiquitin, and β-actin.

**Supplementary Fig. S7.**
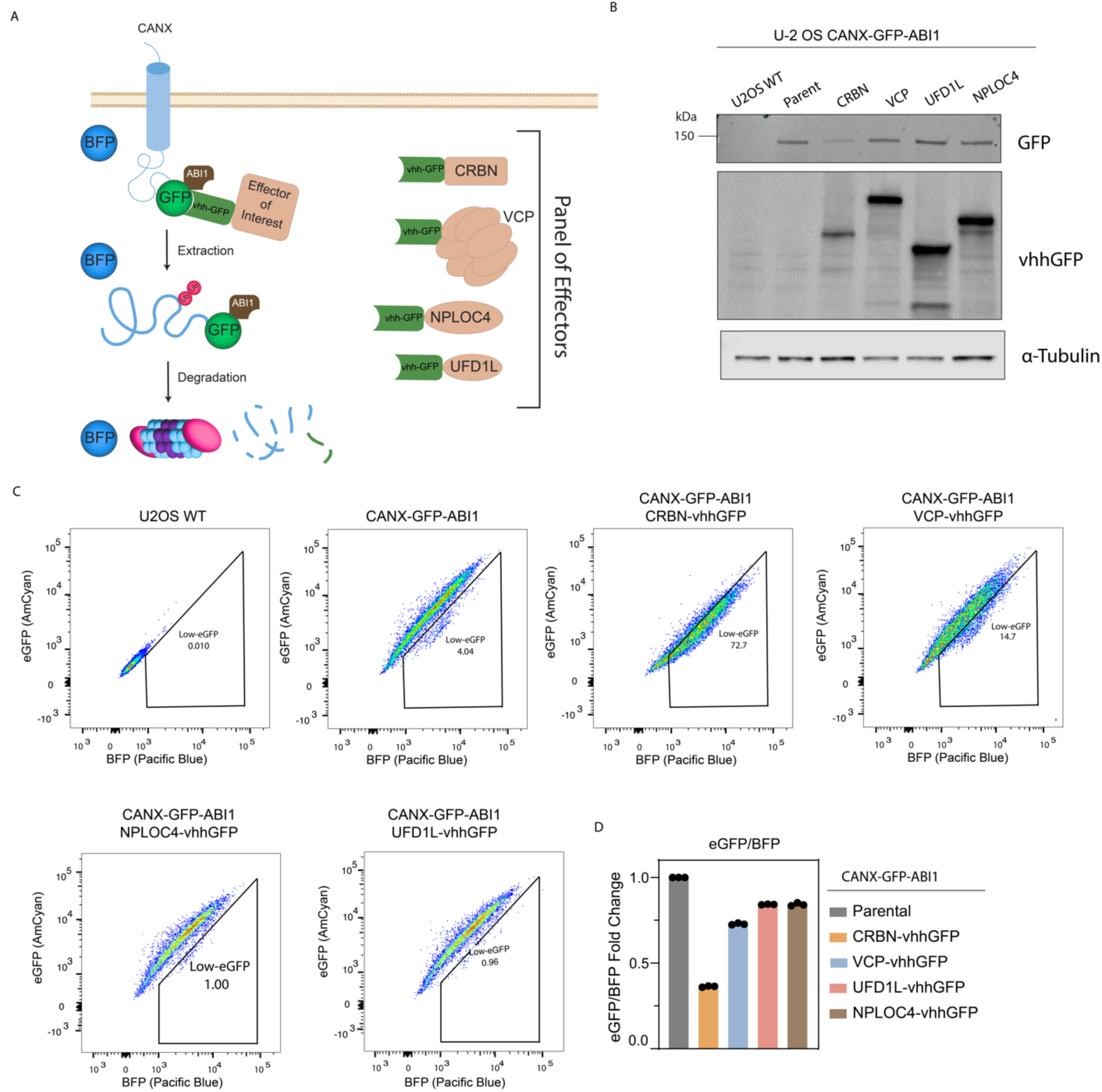
VCP complex recruitment does not induce the degradation of ER-TM proteins. **(A)** Diagram of constitutive induced proximity system with U-2 OS CANX-GFP-ABI1 cells infected with the Effector-vhhGFP CRBN, VCP, NPLOC44, UFD1L. **(B)** Immunoblot of specified cell lines blotting for GFP, vhhGFP, and α-tubulin. **(C)** Scatter plot of GFP (y-axis) and BFP (x-axis) measured using flow cytometry in indicated U-2 OS cell lines expressing constructive induced proximity system and **(D)** quantification of GFP/BFP fold change of Effector-vhhGFP compared to parental control.

**Supplementary Fig. S8.**
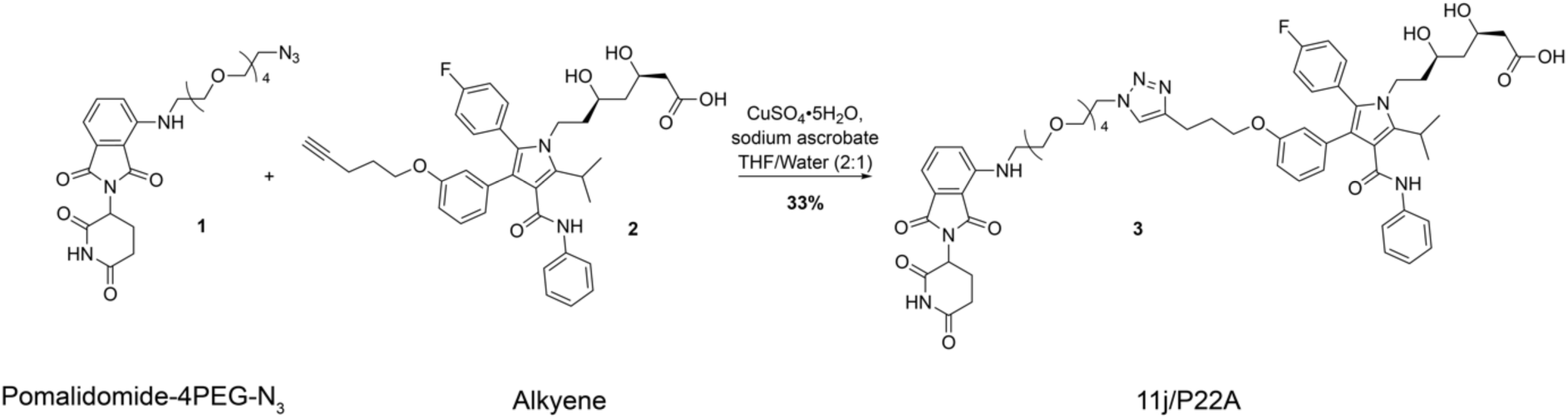
Synthesis of endogenous HMGCR PROTAC, 11j/P22A. 11j/P22A **3** was synthesized from pomalidomide-4PEG-N_3_ **1** and alkyene **2.**

