## Supplemental Table S2 for "Induced ubiquitination bypasses canonical ERAD to drive ER protein degradation"

| Reagent or Resource | Source / Company | Catalog or Reference Number | Additional Information / Use |
|---|---|---|---|
| **Plasmids** | | | |
| pLEX_305-C-dTAG | Addgene | #91798 | Gateway-compatible destination vector |
| pLenti-Cas9-blast | Addgene | #52962 | Cas9 expression vector |
| pMD2.G | Addgene | #12259 | Lentiviral envelope plasmid |
| psPAX2 | Addgene | #12260 | Second-generation packaging plasmid |
| pMDLg/pRRE | Addgene | #12251 | Third-generation packaging plasmid |
| pRSV-Rev | Addgene | #12253 | Third-generation packaging plasmid |
| BFP-KDEL | Addgene | #49150 | ER marker plasmid |
| pLX301-eGFP-ABI1-IRES-BFP | Gift from Taipale Lab | — | Dual-fluorescent construct |
| pLX301-ccDB-vhhGFP | Gift from Taipale Lab | — | Nanobody effector tagging |
| pLX301-CANX-GFP-ABI1-BFP | This study | — | Generated via Gateway cloning |
| pDONR221 | Invitrogen | #11789020 | Gateway entry vector |
| BP Clonase II Enzyme Mix | Invitrogen | #11789020 | Used for BP recombination |
| LR Clonase II Enzyme Mix | Invitrogen | #11791020 | Used for LR recombination |
| ccdB Survival Competent Cells | Invitrogen | #A10460 | For cloning ccdB-containing plasmids |
| GFP-P2A-BFP gBlock | Integrated DNA Technologies (IDT) | — | 5’ atggtcagcaagggggaagagctgttc actggtgtagtccccatacttgtagagcttga cggtgacgtgaacggacataaatttagcgtt tcaggcgagggagagggtgacgcgacat atgggaaactgacactcaagttcatatgtac tactggtaagctccctgtgccctggccgactc tggttaccacgcttacttatggtgtacaatgttt cagtagatatccggaccatatgaagcaaca tgacttctttaaaagtgccatgcctgaagggt atgtacaagaacgcacaattttttttaaagac gatggtaattataagactagggcggaggtta aattcgagggagatacgctggtaaaccgc attgagcttaagggcattgatttcaaagagg acggcaacattttgggacacaaacttgaat |

| | | | |
|---|---|---|---|
| | | | ataactacaatagtcataacgtttacattatggcggacaaacaaaaaaacgggataaaagtgaactttaagattcgccataatatagaggatggaagtgtacaactggcggatcactaccagcagaacactcccattggtgacggtcccgtcttgctgcccgataatcactacttgtccacgcagagtgcgttgagtaaggatccgaatgaaaagagggaccatatggtgctccttgaatttgttaccgcagcgggcatcacgctcggcatggacgagctttacaa 3’ |
| Cytosolic-GFP gBlock | Integrated DNA Technologies (IDT) | — | 5’GGGGACAAGTTTGTACAAAAAAGCAGGCTTCACCatggtgagcaagggcgaggagctgttcaccggggtggtgcccatcctggtcgagctggacggcgacgtaaacggccacaagttcagcgtgtccggcgagggcgagggcgatgccacctacggcaagctgaccctgaagttcatctgcaccaccggcaagctgcccgtgccctggcccaccctcgtgaccaccctgacctacggcgtgcagtgcttcagccgctaccccgaccacatgaagcagcacgacttcttcaagtccgccatgcccgaaggctacgtccaggagcgcaccatcttcttcaaggacgacggcaactacaagacccgcgccgaggtgaagttcgagggcgacaccctggtgaaccgcatcgagctgaagggcatcgacttcaaggaggacggcaacatcctggggcacaagctggagtacaactacaacagccacaacgtctatatcatggccgacaagcagaagaacggcatcaaggtgaacttcaagatccgccacaacatcgaggacggcagcgtgcagctcgccgaccactaccagcagaacacccccatcggcgacggccccgtgctgctgcccgacaaccactacctgagcacccagtccgccctgagcaaagaccccaacgagaagcgcgatcacatggtcctgctggagttcgtgaccgccgccgggatcactctcggcatggacgagctgtacaagCACCCAGCTTTCTTGTACAAAGTGGTCCCC 3’ |
| ER Lumenal-GFP gBlock | Integrated DNA Technologies (IDT) | — | 5’GGGGACAAGTTTGTACAAAAAAGCAGGCTTCATGGATATGCGCGTGCTGGCGCAGCTGCTGGGCCTGCTGCTGCTGTGCTTTCCGGGCGCGCGCTGCGTCAGCAAGGGGGAAGAGCTGTTCACTGGTGTAGTCCCCATACTTG |

| | | | |
|---|---|---|---|
| | | | TAGAGCTTGACGGTGACGTGA<br>ACGGACATAAATTTAGCGTTT<br>CAGGCGAGGGAGAGGGTGAC<br>GCGACATATGGGAAACTGACA<br>CTCAAGTTCATATGTACTACTG<br>GTAAGCTCCCTGTGCCCTGGC<br>CGACTCTGGTTACCACGCTTA<br>CTTATGGTGTACAATGTTTCAG<br>TAGATATCCGGACCATATGAA<br>GCAACATGACTTCTTTAAAAGT<br>GCCATGCCTGAAGGGTATGTA<br>CAAGAACGCACAATTTTTTTTA<br>AAGACGATGGTAATTATAAGA<br>CTAGGGCGGAGGTTAAATTCG<br>AGGGAGATACGCTGGTAAACC<br>GCATTGAGCTTAAGGGCATTG<br>ATTTCAAAGAGGACGGCAACA<br>TTTTGGGACACAAACTTGAATA<br>TAACTACAATAGTCATAACGTT<br>TACATTATGGCGGACAAACAA<br>AAAAACGGGATAAAAGTGAAC<br>TTTAAGATTCGCCATAATATAG<br>AGGATGGAAGTGTACAACTGG<br>CGGATCACTACCAGCAGAACA<br>CTCCCATTGGTGACGGTCCCG<br>TCTTGCTGCCCGATAATCACT<br>ACTTGTCCACGCAGAGTGCGT<br>TGAGTAAGGATCCGAATGAAA<br>AGAGGGACCATATGGTGCTCC<br>TTGAATTTGTTACCGCAGCGG<br>GCATCACGCTCGGCATGGAC<br>GAGCTTTACAAAAAAGATGAA<br>CTGCACCCAGCTTTCTTGTAC<br>AAAGTGGTCCCC3’ |
| HMGCR-GFP gBlock | Integrated DNA Technologies (IDT) | — | 5’GGGGACAAGTTTGTACAAAA<br>AAGCAGGCTTCACCATGTTGT<br>CAAGACTTTTTCGAATGCATG<br>GCCTCTTTGTGGCCTCCCATC<br>CCTGGGAAGTCATAGTGGGG<br>ACAGTGACACTGACCATCTGC<br>ATGATGTCCATGAACATGTTTA<br>CTGGTAACAATAAGATCTGTG<br>GTTGGAATTATGAATGTCCAA<br>AGTTTGAAGAGGATGTTTTGA<br>GCAGTGACATTATAATTCTGA |

| | | | |
|---|---|---|---|
| | | | CAATAACACGATGCATAGCCA<br>TCCTGTATATTTACTTCCAGTT<br>CCAGAATTTACGTCAACTTGG<br>ATCAAAATATATTTTGGGTATT<br>GCTGGCCTTTTCACAATTTTCT<br>CAAGTTTTGTATTCAGTACAGT<br>TGTCATTCACTTCTTAGACAAA<br>GAATTGACAGGCTTGAATGAA<br>GCTTTGCCCTTTTTCCTACTTT<br>TGATTGACCTTTCCAGAGCAA<br>GCACATTAGCAAAGTTTGCCC<br>TCAGTTCCAACTCACAGGATG<br>AAGTAAGGGAAAATATTGCTC<br>GTGGAATGGCAATTTTAGGTC<br>CTACGTTTACCCTCGATGCTC<br>TTGTTGAATGTCTTGTGATTGG<br>AGTTGGTACCATGTCAGGGGT<br>ACGTCAGCTTGAAATTATGTG<br>CTGCTTTGGCTGCATGTCAGT<br>TCTTGCCAACTACTTCGTGTTC<br>ATGACTTTCTTCCCAGCTTGT<br>GTGTCCTTGGTATTAGAGCTT<br>TCTCGGGAAAGCCGCGAGGG<br>TCGTCCAATTTGGCAGCTCAG<br>CCATTTTGCCCGAGTTTTAGA<br>AGAAGAAGAAAATAAGCCGAA<br>TCCTGTAACTCAGAGGGTCAA<br>GATGATTATGTCTCTAGGCTT<br>GGTTCTTGTTCATGCTCACAG<br>TCGCTGGATAGCTGATCCTTC<br>TCCTCAAAACAGTACAGCAGA<br>TACTTCTAAGGTTTCATTAGGA<br>CTGGATGAAAATGTGTCCAAG<br>AGAATTGAACCAAGTGTTTCC<br>CTCTGGCAGTTTTATCTCTCTA<br>AAATGATCAGCATGGATATTG<br>AACAAGTTATTACCCTAAGTTT<br>AGCTCTCCTTCTGGCTGTCAA<br>GTACATCTTCTTTGAACAAACA<br>GAGACAGAATCTCACCCAGCT<br>TTCTTGTACAAAGTGGTCCCC<br>3’ |
| **DNASU Entry Plasmids** | | | |
| CANX | DNASU | HsCD0003<br>9480 | Entry plasmid |

| | | | |
|---|---|---|---|
| LDM (CYP51A1) | DNASU | HsCD00040181 | Entry plasmid |
| EPHX1 | DNASU | HsCD00041470 | Entry plasmid |
| RTN4b | DNASU | HsCD00081743 | Entry plasmid |
| BCAP31 | DNASU | HsCD00719222 | Entry plasmid |
| PKD2 | DNASU | HsCD00082611 | Entry plasmid |
| CRBN | DNASU | HsCD00353201 | Entry plasmid |
| VCP | DNASU | HsCD00829538 | Entry plasmid |
| UFD1L | DNASU | HsCD00081676 | Entry plasmid |
| NPLOC4 | DNASU | HsCD00041106 | Entry plasmid |
| **Primers (UBALER Screen Preparation)** | | | |
| oMCB_1562 | — | — | aggcttggatttctataacttcgtatagcatacattatac |
| oMCB_1563 | — | — | acatgcatggcggtaatacggttatc |
| oMCB_1439 | — | — | caagcagaagacggcatacgagatgcacaaaaggaaactcaccct |
| oMCB_1440-AD002 | — | — | aatgatacggcgaccaccgagatctacacGATCGGAAGAGCACACGTCTGAACTCCAGTCACCGATGTCGACTCGGTGCCACTTTTTC |
| oMCB_1440-AD003 | — | — | aatgatacggcgaccaccgagatctacacGATCGGAAGAGCACACGTCTGAACTCCAGTCACTTAGGCCGACTCGGTGCCACTTTTTC |
| oMCB_1440-AD009 | — | — | aatgatacggcgaccaccgagatctacacGATCGGAAGAGCACACGTCTGAACTCCAGTCACGATCAGCGACTCGGTGCCACTTTTTC |
| oMCB_1440-AD010 | — | — | aatgatacggcgaccaccgagatctacacGATCGGAAGAGCACACGTCTG |

| | | | |
|---|---|---|---|
| | | | AACTCCAGTCACTAGCTTCGA CTCGGTGCCACTTTTTC |
| oMCB_1440-AD022 | — | — | aatgatacggcgaccaccgagatctacac GATCGGAAGAGCACACGTCTG AACTCCAGTCACCGTACGCGA CTCGGTGCCACTTTTTC |
| oMCB_1440-AD025 | — | — | aatgatacggcgaccaccgagatctacac GATCGGAAGAGCACACGTCTG AACTCCAGTCACACTGATCGA CTCGGTGCCACTTTTTC |
| oMCB_1672 | — | — | GCCACTTTTTCAAGTTGATAAC GGACTAGCCTTATTTAAACTTG CTATGCTGTTTCCAGCTTAGC TCTTAAAC |
| **Cell Lines and Culture Media** | | | |
| HEK293, HEK293T, U-2 OS, Huh7 | UC Berkeley Cell Culture Facility | — | Verified mycoplasma-free |
| DMEM (4.5 g/L glucose, L-glutamine, no sodium pyruvate) | Corning | #10-013-CMR | Base medium |
| Fetal Bovine Serum (FBS) | Corning | 35-016-CV | 10% (v/v) |
| Penicillin–Streptomycin | Life Technologies | #15140122 | 1X concentration |
| Blasticidin | Thermo Fisher Scientific | #A1113903 | 4–10 µg/mL |
| TrypLE Express | Gibco | #12605010 | Cell detachment reagent |
| DPBS | Gibco | #1419-144 | Wash buffer |
| Phenol Red-Free Medium | HyClone | #16777-406 | For flow cytometry |
| Lipoprotein-deficient FBS | Kalen Biomedical LLC | #NC9764296 | For Huh7 experiments |
| Hexadimethrine bromide (Polybrene) | Sigma Aldrich | #107689 | 8 µg/mL during transduction |
| **Antibodies** | | | |

| | | | |
|---|---|---|---|
| Anti-VCP | Novus | NB100-1558 | 1:1000 dilution |
| Anti-GFP | Sigma-Aldrich | #SAB1305545 | 1:1000 dilution |
| Anti-HA | Sigma-Aldrich | #H9658 | 1:1000 dilution |
| Anti-α-Tubulin | Cell Signaling Technology | #11H10 | 1:1000 dilution |
| Anti-β-Actin (ACTB) | Sigma-Aldrich | #A5441 | 1:1000 dilution |
| Anti-Ubiquitin | Enzo Life Sciences | #BML-PW0930 | 1:1000 dilution |
| Anti-VHH (rabbit anti-camelid) | GenScript | #A01861 | HRP conjugated |
| IRDye 800CW Goat Anti-Rabbit | LI-COR Biosciences | #926-32211 | Secondary antibody |
| Alexa Fluor 680 Goat Anti-Mouse | Invitrogen | #A21058 | Secondary antibody |
| **Chemicals and Inhibitors** | | | |
| Pierce Protease and Phosphatase Inhibitor Mini Tablets (EDTA-free) | Thermo Scientific | #A32961 | Added to lysis buffers |
| N-Ethylmaleimide (NEM) | Thermo Fisher Scientific | #E1271-1G | 10 mM final |
| Digitonin | Neta Scientific | #SIAL-300410 | 1% in lysis buffer |
| MG132 | Selleck Chemicals | #S2619 | 10 µM proteasome inhibitor |
| Bafilomycin A1 | Sigma-Aldrich | #B1793 | 250 nM lysosome inhibitor |
| CB-5083 | Cayman Chemical | #C814B92 | 5 µM VCP inhibitor |
| NMS-873 | Fisher Scientific | #501015294 | 25 µM allosteric VCP inhibitor |
| MLN4924 | ChemieTek | #CT-M4924 | 500 nM NEDD8-activating enzyme inhibitor |

| | | | |
|---|---|---|---|
| **dTAGv-1** | Tocris Bioscience | #69145 | Degrader compound; 100 nM for cytosolic GFP, 500 nM for ER-TM proteins |
| dTAG-13 | Sigma-Aldrich | #SML2601 | Degrader compound |
| Emetine | Sigma-Aldrich | #E2375-50MG | Translation elongation inhibitor |
| Atorvastatin | Spectrum Chemical | #TCI-A2476 | HMGCR pathway control |
| 11j/P22A | Synthesized in-house | — | HMGCR PROTAC |
| RapiGest SF Surfactant | Waters | #18600809 0 | Proteomics reagent |
| TrypLE Express | Gibco | # 12-605-010 | Protease for digestion |
| $CuSO_4 \cdot 5H_2O$ | — | — | Catalyst for click chemistry |
| Sodium ascorbate | — | — | Reducing agent for click reaction |
| THF | — | — | Solvent |
| Dimethyl sulfoxide (DMSO) | Sigma-Aldrich | #472301 | Solvent |
| **Kits and Enzymes** | | | |
| RNeasy Mini Kit | Qiagen | #74104 | RNA isolation |
| Maxima First Strand cDNA Synthesis Kit | Thermo Fisher Scientific | #K1641 | cDNA synthesis |
| TaqMan Gene Expression Assays | Applied Biosystems | #4453320 #30034593 8 #4453320 #30034717 0 | qPCR probe sets |
| TaqMan Universal PCR Master Mix II | Applied Biosystems | — | qPCR master mix |
| QIAquick Gel Extraction Kit | Qiagen | #28704 | DNA purification |
| QIAamp DNA Blood Midi Kit | Qiagen | — | gDNA extraction |
| Herculase II Fusion DNA Polymerase | Agilent | #600679 | Library PCR amplification |

| | | | |
|---|---|---|---|
| GFP-Trap Magnetic Agarose | Proteintech (ChromoTek ) | #gtma | Immunoprecipitation |
| **Consumables and Equipment** | | | |
| 4–20% Mini-PROTEAN TGX Gels | Bio-Rad | — | SDS-PAGE separation |
| Nitrocellulose Membranes | Bio-Rad | — | Blotting |
| PVDF Membranes | Bio-Rad | — | For ubiquitin blots |
| Nunc Lab-Tek II Chambered Coverglass | Thermo Fisher Scientific | #155360 | Live-cell imaging |
| Zeiss Axio Observer 7 Microscope | Zeiss | — | Live imaging |
| LI-COR Odyssey Imager | LI-COR Biosciences | — | Fluorescent blot imaging |
| ChemiDoc MP Imaging System | Bio-Rad | — | HRP blot detection |
| BD LSR Fortessa | BD Biosciences | — | Flow cytometer |
| BD FACS Aria Fusion | BD Biosciences | — | Cell sorter |
| FlowJo v10 | BD Biosciences | — | Flow cytometry analysis |
| GraphPad Prism v8–10 | GraphPad | — | Statistical analysis |
| Bowtie2 software | — | — | sgRNA alignment |
| casTLE framework | — | — | Screen hit scoring |
